# Reovirus uses temporospatial compartmentalization to orchestrate core versus outercapsid assembly

**DOI:** 10.1101/2022.06.06.494974

**Authors:** Justine Kniert, Theodore dos Santos, Heather E. Eaton, Woo Jung Cho, Greg Plummer, Maya Shmulevitz

## Abstract

*Reoviridae* virus family members, such as mammalian orthoreovirus (reovirus), encounter a unique challenge during replication. To hide the dsRNA from host recognition, the genome remains encapsidated in transcriptionally active proteinaceous core capsids that transcribe and release +RNA. *De novo* +RNAs and core proteins must repeatedly assemble into new progeny cores in order to logarithmically amplify replication. Reoviruses also produce outercapsid (OC) proteins µ1, σ3 and σ1 that assemble onto cores to create highly stable infectious full virions. Current models of reovirus replication position amplification of transcriptionally-active cores and assembly of infectious virions in shared factories, but we hypothesized that since assembly of OC proteins would halt core amplification, OC assembly is somehow regulated. Using kinetic analysis of virus +RNA, core and OC proteins, core assembly and whole virus assembly, assembly of OC proteins was found to be temporally delayed. All viral RNAs and proteins were made simultaneously, eliminating the possibility that delayed OC RNAs or proteins account for delayed OC assembly. High resolution fluorescence and electron microscopy revealed that core amplification occurred early during infection at peripheral core-only factories, while all OC proteins associated with lipid droplets (LDs) that coalesced near the nucleus in a µ1–dependent manner. Core-only factories transitioned towards the nucleus despite cycloheximide-mediated halting of new protein expression, while new core-only factories developed in the periphery. As infection progressed, OC assembly occurred at LD-and nuclear-proximal factories. Silencing of OC µ1 expression with siRNAs led to large factories that remained further from the nucleus, implicating µ1 in the transition to perinuclear factories. Moreover, late during infection, +RNA pools largely contributed to the production of *de-novo* viral proteins and fully-assembled infectious viruses. Altogether the results suggest an advanced model of reovirus replication with spatiotemporal segregation of core amplification, OC complexes and fully assembled virions.

**NON-TECHNICAL AUTHOR SUMMARY:** It is important to understand how viruses replicate and assemble to discover antiviral therapies and to modify viruses for applications like gene therapy or cancer therapy. Reovirus is a harmless virus being tested as a cancer therapy. Reovirus has two coats of proteins, an inner coat and an outer coat. To replicate, reovirus particles need only the inner coat, but to become infectious they require the outer coat. Strangely, inner and outer coat proteins are all made by the virus at once, so it was unknown what determines whether newly made viruses will contain just the inner coat to continue to replicate, or both coats to transmit to new hosts. Our experiments reveal that the inner coat proteins are located in a different area of an infected cell versus the outer coat proteins. The location therefore determines if the newly made viruses contain just the inner coat versus both coats. Reoviruses have evolved extravagant mechanisms to be able to efficiently take on the best composition required for replication and transmission.

## INTRODUCTION

The *Reoviridae* family encompasses viruses that impact the well-being of many domains of life, including agricultural pathogens such as Rice dwarf virus, livestock pathogens such as bluetongue virus and African horse sickness virus, and aquaculture pathogens such as grass carp reovirus^1, 2^. In humans, rotavirus is a major cause of severe childhood diarrhoea, but thankfully rotavirus vaccines have greatly reduced childhood mortality especially in developing countries. There are also rare emerging zoonotic pathogens in the *Reoviridae* family such as encephalitis-associated Banna virus, Kadipiro virus and Liao ning virus. One of the best-studied members of the *Reoviridae* family is mammalian orthoreovirus (reovirus). Reovirus infects the majority of humans by adolescence, as well as a wide variety of mammals though the fecal-oral route, but causes minimal-to-no disease symptoms^3–9^. In addition to replicating harmlessly in the enteric tract^10–15^, reovirus can replicate in transformed cells of tumors, kill cancer cells and augment anti-tumor immunity; accordingly, reovirus is undergoing clinical testing as a cancer therapy^16–21^. Importantly, reovirus provides a safe model system to study the *Reoviridae* family.

All *Reoviridae* share a segmented dsRNA genome that must be shielded from host innate detection receptors, and therefore the fundamental unit of replication for all *Reoviridae* is a core capsid that surrounds the genome and never disassembles^1, 2^. Inside the core, a viral polymerase and co-factors synthesize positive-sense RNAs from dsRNA templates, which are capped by a viral capping enzyme following release from the cores into the cellular cytoplasm. Viral proteins are synthesized by host translation machinery. Core proteins and positive-sense RNAs are assembled into new progeny cores that synthesize negative-sense RNA within the core and amplify the replication process. Some *Reoviridae* exist only as cores, such as members of plant- and insect-infecting oryzavirus, cypovirus and Dinovernavirus genera. But most *Reoviridae* have evolved an outercapsid (OC) that surrounds the core when the virus is extracellular. The OC imparts added stability to the viruses, allowing them to persist in bodies of water for months to years. The OC proteins also mediate attachment to cells and confer the ability to transverse cellular membranes during entry. In summary, the core is the replication unit, while the whole virus, consisting of the core and OC proteins, is the infectious unit of OC-containing *Reoviridae*.

Many studies on reoviruses have revealed the processes of virus entry into cells and OC disassembly (uncoating). Specifically for reovirus, the outermost OC protein σ3 (Figure 1A) is degraded by intestinal or lysosomal proteases. The intermediate OC protein µ1 is cleaved to a hydrophobic δ fragment that mediates membrane penetration to deliver the replication-component core to the cellular cytoplasm. What is poorly understood is how the assembly of core versus OC proteins of reovirus is orchestrated. It has long been recognized that reovirus replicates within neo-organelles/factories formed by the non-structural protein μNS, which recruits ssRNA-binding non-structural protein σNS, tubulin-binding core μ2 protein, core proteins σ2, λ1, λ2 and λ3 and a variety of host components, such as endoplasmic reticulum remnants and cytoskeletal elements^22–25^. But the central dogma depicts reovirus core replication and OC assembly occurring in the same factories. This current model presents a critical conundrum: once the OC is assembled onto cores, the cores become transcriptionally inactive and cannot amplify replication. How then do cores persist as cores to amplify replication? This replication-versus-OC assembly conundrum raises the hypothesis that core versus full assembly is orchestrated, such that OC assembly is in some way delayed to permit core amplification without full assembly.

**Figure 1.**
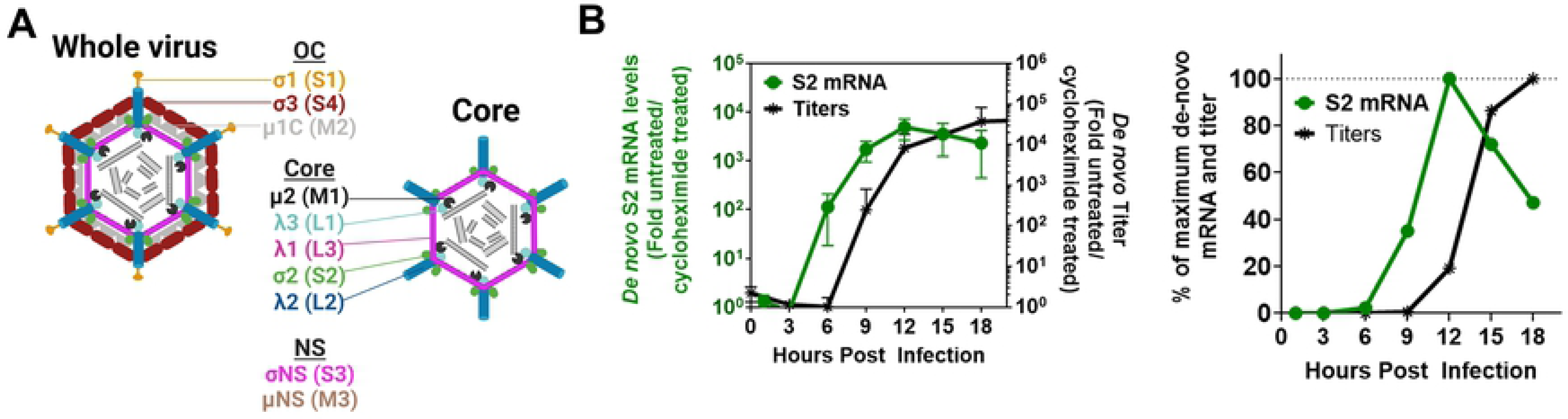
Reovirus infectious virus production is delayed relative to core amplification. **(A)** Diagram of reovirus whole virion and core indicating respective capsid proteins. In brackets are the genome segments encoding each protein. Non-structural proteins and their genome segments are also listed. Created with Biorender.com. **(B)** L929 cells were infected with reovirus at an MOI of 3 with or without cycloheximide (100μg/mL) and samples were collected every 3 hours to measure S2 mRNA levels and titers. **(B, Left)** *De novo* S2 mRNA levels by RT-qPCR and virus titers at each time point, calculated as fold-increase in cycloheximide-untreated versus cycloheximide-treated samples for each timepoint, n=4. **(B, Right)** *De novo* mRNA and virus titers calculated as % of maximum achieved at 12 hpi for mRNA and 18 hpi for titers.

In a quest to solve the replication-versus-OC assembly conundrum, we conducted a comprehensive kinetic analysis of virus RNA, protein and whole-virus production. While core replication, represented by RNA and protein logarithmic amplification, occurred between 6-12 hours post infection (hpi), assembly of OCs to generate infectious virions was logarithmically escalated between 9-15 hpi; this suggested a 3-hour delay between core and OC assembly. Moreover, unlike previously surmised, the 10 reovirus mRNAs and the major core and OC proteins had similar rates of synthesis, indicating that the delayed assembly of the OC was not caused by deferred expression of OC RNAs or proteins. To determine if core and OC proteins may be spatially compartmentalized, we generated previously-unavailable antibodies towards core proteins and conduced immunofluorescence microscopy of virus-infected and virus protein-transfected cells. Surprisingly, we discovered 3 distinct localizations for reovirus proteins. First, there were factories at the periphery of infected cells that stained with core-but not OC proteins (“core-only” factories). Second, as previously described^26^, the OC protein µ1 predominated on lipid droplets (LDs) during virus infection; but herein we discovered that all three OC proteins presided on LDs in a µ1-dependent manner. Third, there were perinuclear factories positive for both core and OC proteins (“core-and-OC factories”). These findings indicate that core and OC proteins are spatially segregated. Electron microscopy confirmed the existence of cores in core-only factories, and fully-assembled viruses in perinuclear factories. Moreover, core-only factories were positive for *de novo* transcription, indicating that core amplification was occurring. Temporally, core-only factories were predominant at 8 hpi, while perinuclear core-and-OC factories expanded at 12 hpi onwards, coinciding with the kinetic analysis of *de novo* RNA versus infectious virus production. Importantly, siRNA-mediated silencing of OC µ1 led to factories that were further from the nucleus and larger, suggesting that LD-associated OC protein localization guides perinuclear localization of factories. Altogether, our study reveals that the temporospatial segregation of reovirus OC proteins at LDs enables uninterrupted amplification of cores in core-only factories, and that as core-only factories move towards the LDs and perinuclear space, full assembly to infectious virions dominates.

## RESULTS

### Reovirus infectious virus production is delayed relative to core amplification

During infection, reovirus exists either as a transcriptionally-active but non-infectious core particle or as a transcriptionally-inert but infectious whole virion consisting of cores coated by OC proteins (Figure 1A).

Assembly of OC proteins on the core particle halts positive RNA synthesis, and therefore terminates further amplification of replication^27^. As such, we hypothesized that in order to establish a productive infection, reovirus cores must first accumulate without OC assembly in order to amplify replication.

To examine if there was a delay in OC assembly relative to core amplification, a kinetic analysis of core-derived mRNA synthesis versus titers of fully-assembled viruses was performed. Tumorigenic L929 mouse fibroblasts, the most commonly used cell line for reovirus studies, were infected with reovirus strain T3D^PL^ (herein referred to as reovirus) at an MOI of 3. Lysates were collected at 3-hour increments for analysis of S2 positive-sense single-stranded (+ssRNA) levels as a measure of core amplification, versus infectious virus titers as a measure of OC assembly. Importantly, infections were performed in the absence or presence of cycloheximide. Cycloheximide halts protein translation and as such, provides a measure of input core transcription and background infectious virions. Bonafide *de novo* RNA and infectious virus levels were achieved by calculating the fold change of RNA and infectious titers for cycloheximide-untreated versus cycloheximide-treated matched samples. Two unique S2-specific primer sets were used to ensure accuracy. Moreover, during purification of RNA from infected cells, RNA purification columns that do not bind dsRNA were used to focus exclusively on +ssRNA and exclude genomic dsRNA. Reovirus *de novo* S2 +ssRNA levels logarithmically escalated between 3-12 hpi (Figure 1B, left). Conversely, reovirus *de novo* infectious virus titers logarithmically escalated between 6-15 hpi, suggesting a 3-hour delay in OC assembly relative to core amplification. When plotted as percent of maximum levels achieved (Figure 1B, right), maximum *de novo* +ssRNA was achieved at 12 hours while maximum infectious titers at 18 hours. At 9 hpi, 40% of maximal *de novo* +ssRNA was achieved, yet no new infectious viruses produced. Interestingly, between 12-18 hpi, *de novo* +ssRNA levels decreased relative to maximum levels, likely reflecting the packaging and conversion to dsRNAs. Overall, results of the kinetic analysis suggested that there was an ∼3-hour delay in OC assembly relative to core amplification.

### Core and outercapsid RNAs and proteins are expressed at similar kinetics but assembled at distinct kinetics

One possible mechanism for delayed OC assembly could be delayed synthesis of OC-coding mRNAs or proteins. In the 1970s, several manuscripts suggested that reovirus transcription is regulated^28–31^. Specifically, the authors separated radiolabelled and hybridized products of reovirus ssRNA and dsRNA by electrophoresis on polyacrylamide gel columns and concluded that M1, L2 and L3 mRNA synthesis requires new protein expression, and that not all viral transcripts and proteins are generated at the same time. Given the advent of sensitive RT-qPCR assays, we revisited this dogma by measuring all reovirus +ssRNAs over the course of reovirus replication. Two distinct primer sets were used for each of the 10 reovirus RNAs, and fold-change over cycloheximide treatment overcame differences among primer efficiencies and limits of detection (Supplementary Figure S1).

There were no significant differences in levels of *de novo* synthesized +ssRNA among the 10 reovirus genome segments when absolute levels were compared at each time point (Figure 2A, left). In either case, there were no delays in OC-coding mRNAs relative to core-coding mRNAs that could explain the delayed OC assembly.

**Figure 2.**
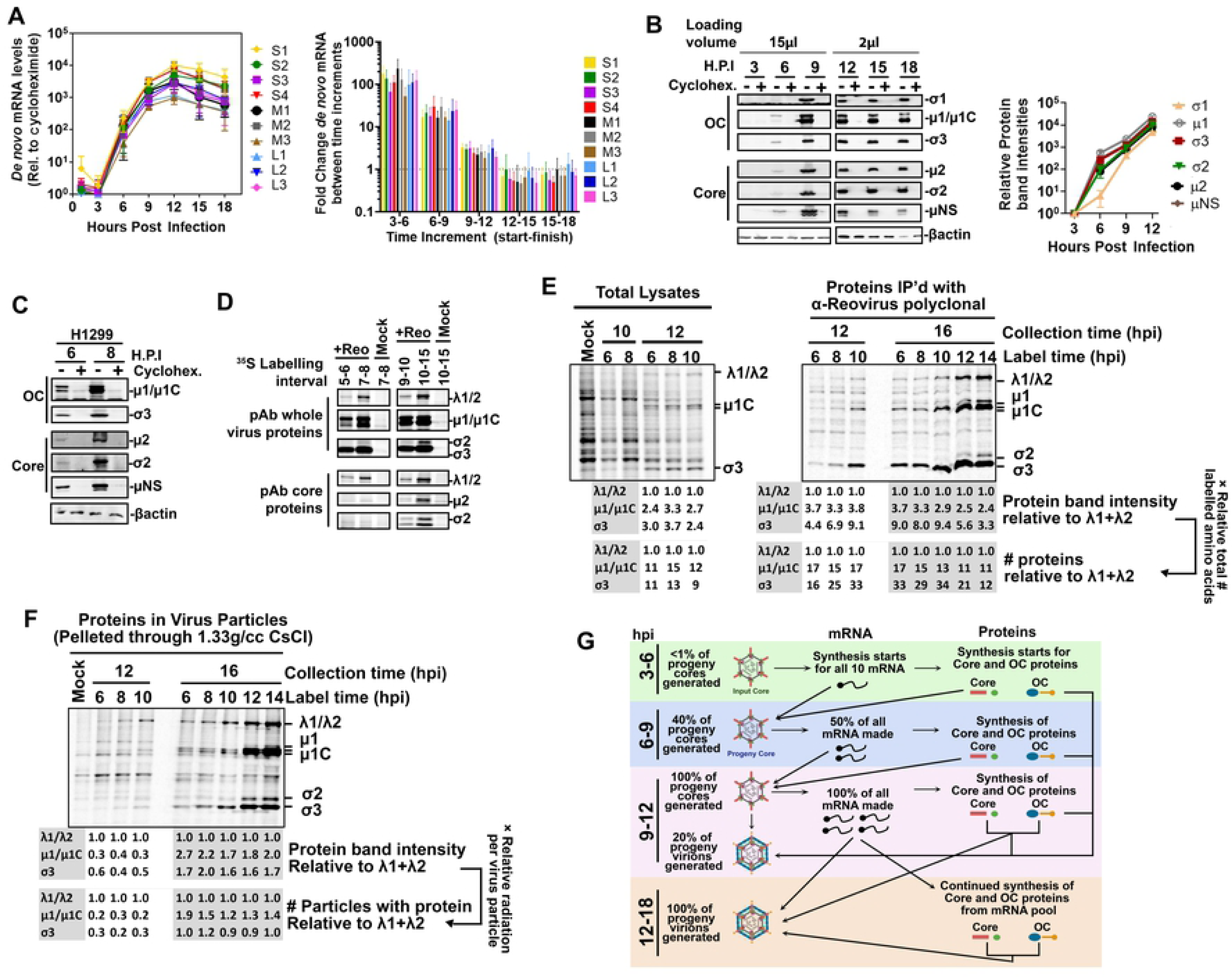
Reovirus RNAs and proteins are generated simultaneously. **(A-B)** L929 cells were infected with reovirus at an MOI of 3 with or without cycloheximide (100μg/mL) and samples were collected every 3 hours to measure virus mRNA and protein levels. **(A, Left)** *De novo* mRNA levels by RT-qPCR at each time point for the indicated reovirus transcripts, calculated as fold-increase in cycloheximide-treated over mock-treated samples for each timepoint, n=4. **(A, Right)** Fold increase in *de-novo* virus mRNA among the time intervals indicated, n=4. **(B, Left)** Representative blots show *de novo* virus protein levels by SDS-PAGE and Western blot analysis using polyclonal antibodies raised against whole virus (detects µ1c and σ3), µ2, σ2, or µNS, or the σ1 N-terminal domain as indicated. **(B, Right)** Combined protein levels for n=4. **(C-D)** H1299 cells were infected with reovirus at an MOI of 3. **(C)** Similar to B but in H1299 cells. **(D)** H1299 cells infected with reovirus radio-labelled with S^35^-methionine/cysteine for the indicated time increments and then immediately lysed. Lysates were subjected to immunoprecipitation with either polyclonal antibodies from rabbits exposed to whole reovirus (pAb whole virus proteins) that predominantly recognize OC µ1c and σ3 but also core proteins λ1/2, or polyclonal antibodies from rabbits exposed to reovirus cores (pAb core proteins) that predominantly recognize core proteins σ2, µ2 and λ1/2. Mock infected cells were used as negative controls. **(E-F)** H1299 cells infected with reovirus (or mock) were pulse-labelled with S^35^-methionine/cysteine at the indicated time (label time) for 30 minutes and then chased with normal media until the indicated collection time. **(E, Left)** Whole lysates show pulse-labelled virus and host proteins. **(E, Right)** Whole lysates were subjected to immunoprecipitation with polyclonal anti-reovirus antibodies prior to electrophoresis. **(F)** Whole lysates were subjected to high-speed ultracentrifugation through 1.33g/cc CsCl to pellet reovirus cores and fully-assembled viruses. Minimal host protein contamination is demonstrated by the mock control. **(G)** Using values from figures 1 and 2, summary of *de-novo* mRNA, proteins, progeny cores, and progeny infectious viruses is presented during the course of one reovirus round of replication. Statistical analysis by ANOVA, **** p<0.0001, *** p<0.001, **p<0.05, ns > 0.05.

To assess if OC protein expression was delayed relative to core protein expression, antibodies had to be generated towards core proteins since already-available antibodies are exclusively focused on reovirus OC or non-structural proteins. Rabbit polyclonal antibodies were generated against whole infectious reovirus particles; these antibodies recognize OC protein µ1 (and the µ1C product generated when µ1C associates with σ3^32^), OC protein σ3 and the core protein λ2 (Supplementary Figure S2A and S2B). Core particles were also generated by *in vitro* digestion of viruses with chymotrypsin^33^, purified and used to generate rabbit polyclonal antibodies towards core proteins; these antibodies recognize core proteins σ2, λ1, and λ2 (Supplementary Figure S2A and S2B). Rabbit antibodies were also generated towards bacterially-expressed and purified core proteins µ2 and σ2 and the non-structural protein µNS. L929s were infected at an MOI of 3 for 3-18h and the antibodies were used to detect reovirus core and OC proteins over the course of infection by Western blot analysis (Figure 2B, left). While it is not accurate to compare the band intensities among different proteins due to variation in antibody affinities, the time course analysis permitted a kinetic comparison of each protein accumulation over time (Figure 2B, right).

OC proteins µ1/µ1C and σ3 had similar expression kinetics to core proteins σ2 and µ2 and to the non-structural protein µNS. The only protein with lower expression was σ1, which is the protruding cell attachment protein at reovirus vertices. The S1 mRNA that encodes σ1 also contains an internal reading frame that encodes a distinct non-structural protein σ1s^34^. The translational start sequence of σ1 is weak (Supplementary Figure S3) to permit ribosome bypass to the σ1s start codon. The lower expression of σ1 is well tolerated by reovirus because while 600 copies of OC µ1 and σ3 are required per virion, each virion only assembles up to 36 copies of σ1^35^. These results suggest that core and OC proteins are generated simultaneously.

Expression of reovirus proteins was further validated in the H1299 human lung carcinoma cell line, which is equally permissive to reovirus as L929 cells. Similar to L929 cells, both OC and core proteins were expressed at early time points (6 and 8 hpi) (Figure 2C). Western blot analysis monitors the accumulation of protein levels but does not indicate the precise expression at a given timepoint, and also becomes saturated beyond 12hpi due to low linear range. As a complementary approach to monitor virus protein expression kinetics, H1299 cells were infected at an MOI of 3 with reovirus and subjected to pulse-chase analysis by the addition of ^35^S-methionine for 1 hour at different timepoints, followed by lysis and immunoprecipitation for OC proteins using anti-reovirus or anti-core polyclonal antibodies (Figure 2D). While it is not accurate to compare among virus proteins because of variation in antibody affinities, the pulse-chase analysis demonstrated very strong expression of OC proteins at early time points. OC proteins µ1/µ1C and σ3 have equal-or-fewer methionine and cysteine residues relative to the core proteins (Supplementary Figure S3) and therefore the high intensity of ^35^S-methionine-labelled OC proteins is not attributed to a higher labelling opportunity.

Overall, OC RNAs and proteins seem to accumulate early during infection, yet our kinetic analysis (Figure 1) indicated minimal assembly of OC on cores. Accordingly, the delayed OC assembly is unlikely to be a consequence of delayed OC RNA or protein synthesis.

Radioactive labelling provided an opportunity to further validate the herein-proposed delay of OC protein assembly. Cells infected at an MOI of 2 were subjected to ^35^S-methionine pulse labelling for 30 minutes at various timepoints, but importantly, lysates were also collected at 10, 12 or 16 hours; this experiment was designed to indicate not only when proteins were synthesized, but when they become incorporated into new virions. Lysates were then assessed for total protein expression (Figure 2E) versus assembly (Figure 2F). In total lysates collected at 10 and 12 hpi, reovirus core and OC proteins produced at 6, 8 or 10hpi became detectable over cellular proteins at 12hpi. From total lysates, it is clear that OC proteins µ1/µ1C and σ3 are more abundant than core λ1 and λ2 proteins (Figure 2E, left). If considering the relative total numbers of methionine/cysteines available for labelling in these proteins (Supplementary Figure S3), there is ≥10-fold higher expression of these OC µ1/µ1C and σ3 proteins relative to core λ1 and λ2. Given similar Kozak scores (Supplementary Figure S3), the higher expression of these OC proteins might reflect their smaller size or other yet-to-be discovered mechanisms for differential expression. Nevertheless, expression of OC proteins was not limiting early during infection. Moreover, immunoprecipitation with polyclonal anti-reovirus antibodies also indicated high OC µ1/µ1C and σ3 expression throughout infection (Figure 2E, right). Surprisingly, despite that mRNA production declined after 12hpi (Figure 2A), *de-novo* protein synthesis remained high at 12 and 14hpi (Figure 2E, right); this suggests that the pool of mRNA produced over the first 12 hours amply supports continued virus protein synthesis beyond 12 hpi. Most importantly, the ^35^S-Metionine-labelled lysates were subjected to high-speed centrifugation through 1.33g/cc cesium chloride to pellet cores and whole viruses that have buoyant densities of 1.43 and 1.36 g/cc respectively.

Particles were then assessed for relative levels of OC proteins µ1/µ1C and σ3 versus core λ1 and λ2 proteins based on band intensity or band-intensity calibrated to relative radiation per virus particle (calculated by number of methionines/cysteines and relative copies of each protein per particles, Supplementary Figure 3). Virus particles assembled at 12 hpi only had ∼25% of the OC composition expected for fully-assembled virions and therefore ∼75% of particles were cores (Figure 2F). Conversely at 16hpi, particles were mostly whole assembled viruses based on composition of core and OC proteins. The core-versus-whole virus composition in Figure 2F corresponds strongly with the relative mRNA versus titer kinetics in Figure 1B, and indicates that OC proteins expressed in the first 10 hours post-infection are minimally assembled onto progeny cores in the first 12 hours of infection.

Altogether, the kinetic analysis revealed a new detailed timeline of the reovirus replication cycle (Figure 2G). In the first 3-6 hours post-infection, reovirus input cores generate all 10 mRNAs equally, and both core and OC proteins. Between 6-9 hours, 40% of maximal progeny cores are assembled without OC proteins, and these progeny cores synthesize 50% of maximal mRNA, and both core and OC expression continues. Between 9 and 12hpi, OC proteins begin to assemble onto ∼25% of progeny cores, but the remaining progeny cores achieve the maximal production of mRNAs. Beyond 12 hpi, +RNAs are assembled into new virions to a greater extent than they are produced, resulting in decreased overall +RNA levels. Virus OC and core proteins continue to be expressed from the pre-existing +RNA pools, and pools of +RNA, core, and OC proteins assembly primarily into infectious virions. Neither RNA or protein abundance can explain why OC assembly onto cores is delayed prior to 12hpi.

### Core and outercapsid proteins are spatially compartmentalized during infection

Another possible mechanism to delay OC assembly could be compartmentalization or segregation of core versus OC proteins in the cells. Previous studies found that reovirus replicates in factories formed by non-structural proteins μNS and σNS, the core μ2 protein, and a variety of host components such as endoplasmic reticulum remnants and cytoskeletal elements^22–24^. Not only do these factories provide protection from cellular innate immune responses, but they also serve as sites for the accumulation of viral particles. A previous study also discovered that µ1/µ1C localized to LDs and while the relationship to virus assembly was not deduced, the authors demonstrated a strong relationship between LD-associated µ1/µ1C and cell death^26^. We questioned if viral factories were homogenous throughout the cell, or if they demonstrated phenotypic differences with respect to core versus OC protein composition.

Spinning disk confocal immunofluorescence microscopy was used to examine if core versus OC proteins were spatially compartmentalized. H1299 cells were used for these studies because they are large and flat, permitting easy detection of fluorescence segregation versus overlap. H1299 cells were infected at an MOI of 3, paraformaldehyde fixed at 14 hpi and subjected to immunofluorescence staining with anti-core polyclonal antibodies, monoclonal antibodies towards OC proteins µ1/µ1C, σ1, or σ3^36^, monoclonal antibodies towards σNS^36^, polyclonal antibodies towards µNS, and/or BODIPY dye for LD detection (Figure 3). When comparing staining for core versus OC proteins µ1/µ1C, σ1 or σ3 (Figure 3A), small foci at the periphery of cells were positive for core-but not OC proteins; these foci are herein referred to as “core-only” regions. In the mid- and perinuclear regions of the cell, larger foci co-stained for core and OC proteins. 3D images created from Z-stacks clearly distinguished core-only peripheral foci from core+σ3-positive regions near the nucleus (Figure 3B, Supplemental Video 1). The 3D images also revealed that some σ3 resided at structures resembling LDs. In 3D images of infected cells stained for core and µ1/µ1C (Figure 3C), µ1/µ1C was exclusively detected on LDs positive for fatty-acid binding BODIPY. The absence of µ1/µ1C at perinuclear regions where σ1 and σ3 reside is likely explained by the fact that when virions are fully assembled, µ1/µ1C is hidden under σ3 and not exposed to antibodies. Finally, the factory-forming non-structural protein µNS phenocopied core staining, with distinct peripheral factories relative to OC capsid proteins (Figure 3D). Moreover, the core and factory-forming non-structural protein σNS exhibited strong overlap with anti-core (Figure 3E), indicating that factory-forming non-structural proteins were in core-only foci. Similar results were obtained in human T47D breast cancer cells, another reovirus-permissive cell line conducive to immunofluorescence analysis (Supplemental Figure S4).

**Figure 3.**
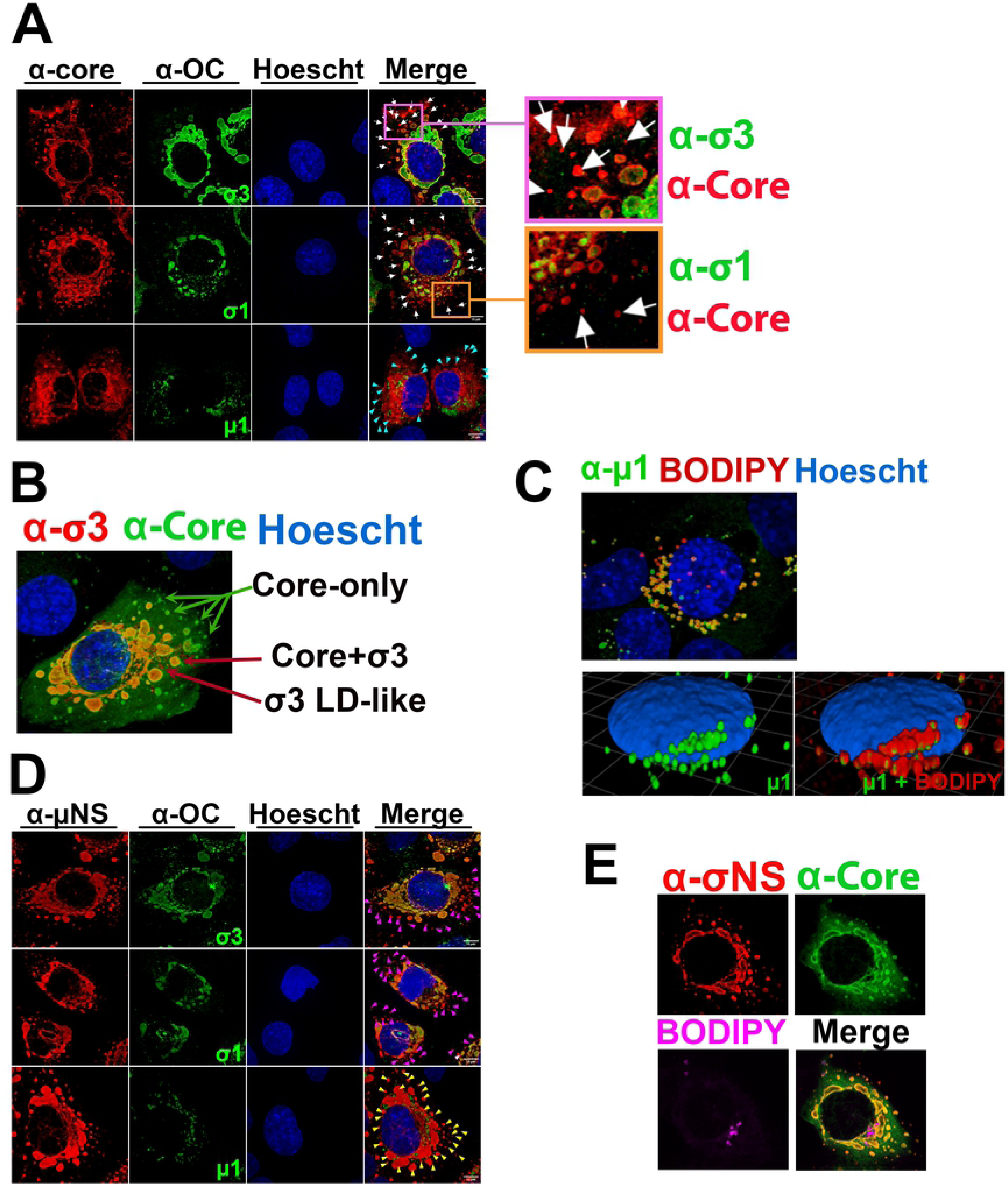
Reovirus proteins are spatially compartmentalized. **(A)** H1299 cells were infected with reovirus at an MOI of 3 before fixation at 14-20 hpi. Immunofluorescence staining was conducted with antibodies specific to OC proteins indicated in green. The OC proteins were detected with secondary antibodies conjugated to Alexa 488 (pseudo colored green). Co-immunofluorescence in the same cells was conducted using polyclonal rabbit antibodies raised against reovirus cores (α-Core) detected with secondary antibodies conjugated to Alexa 647 (red). In the merged images and their corresponding zoomed-in regions, white arrows show example regions of core-only staining, while cyan arrows indicate core-positive but μl-negative regions. Similar results were also obtained with monoclonal 10C1 and 4F2 for σ3, 8H6 for µ1, and rabbit σ1-specific polyclonal serum (data not shown). **(B)** Represented 3D images created from Z-stacks of reovirus infected H1299 cells probed against σ3 (10C1, red), core (α-Core, green), and nuclei (Hoechst dye, blue). **(C)** Represented 3D images created from Z-stacks of reovirus infected H1299 cells probed against μl (10F6, green), LDs (BODIPY dye, red), and nuclei (Hoechst dye, blue). (bottom) The BODIPY channel showing LDs is toggled off (left) and on (right). **(D)** Similar to (A) but rabbit polyclonal antibodies to non-structural protein μNS (α-μNS, red) were used instead of α-Core for co-immunofluorescence with antibodies directed towards the OC proteins indicated [deleted words here]. Magenta arrows indicate regions that are μNS-positive but OC-negative, and yellow arrows indicated μNS-positive but μl-negative regions. **(E)** Represented 3D images created from Z-stacks of reovirus infected H1299 cells probed against σNS (mouse monoclonal 2A9, red), core (α-Core, green), and LDs (BODIPY dye, magenta). **(A-E)** All images were acquired using spinning disk confocal microscopy and analyzed with Volocity software. Images are representative of at least four images captured for each condition from seven biologically independent experiments.

Altogether, reovirus proteins seemed to segregate into 3 distinct regions of the cell. First, core-only regions in the periphery of cells consist of core proteins and non-structural proteins, but not OC proteins. Second, OC proteins µ1 and σ3 but not core proteins localize at LDs. Third, perinuclear regions contain both core and OC proteins σ3 and σ1.

### Outercapsid proteins localize to lipid droplets in transfected cells

Monitoring virus protein localization during infection in Figure 3 came with two challenges. First, it was difficult to ascertain if the spatial segregation of reovirus core- and OC proteins was an inherent property of the virus proteins, or rather a consequence of infection and virus amplification; in other words, would these proteins localize to respective regions without virus infection? Secondly, the fluorescently-intense high accumulation of OC σ3 at the perinuclear regions made it challenging to establish if there was bonafide σ3 localization also at LDs; in other words, while µ1/µ1C is easily seen on LDs because it is hidden in fully-assembled virions, σ3 at LDs could be overpowered by the much-stronger signals at the perinuclear regions. Therefore, to conclusively determine where OC proteins localize in absence of infection, H1299 cells were transfected with eukaryotic plasmids that constitutively express the non-structural protein μNS required for pseudo-factory formation, along with OC proteins σ1 and σ3, in the presence or absence of μ1. Cells were stained for OC proteins in combination with BODIPY to visualize LDs (Figure 4A, Supplementary Figure S5). Along with μ1, both σ1 and σ3 staining revealed ring-like structures surrounding the outer surface of LDs. LD localization of σ1 and σ3 was contingent on the presence of μ1, indicating that μ1 facilitates localization of OC σ1 and σ3 at LDs.

**Figure 4.**
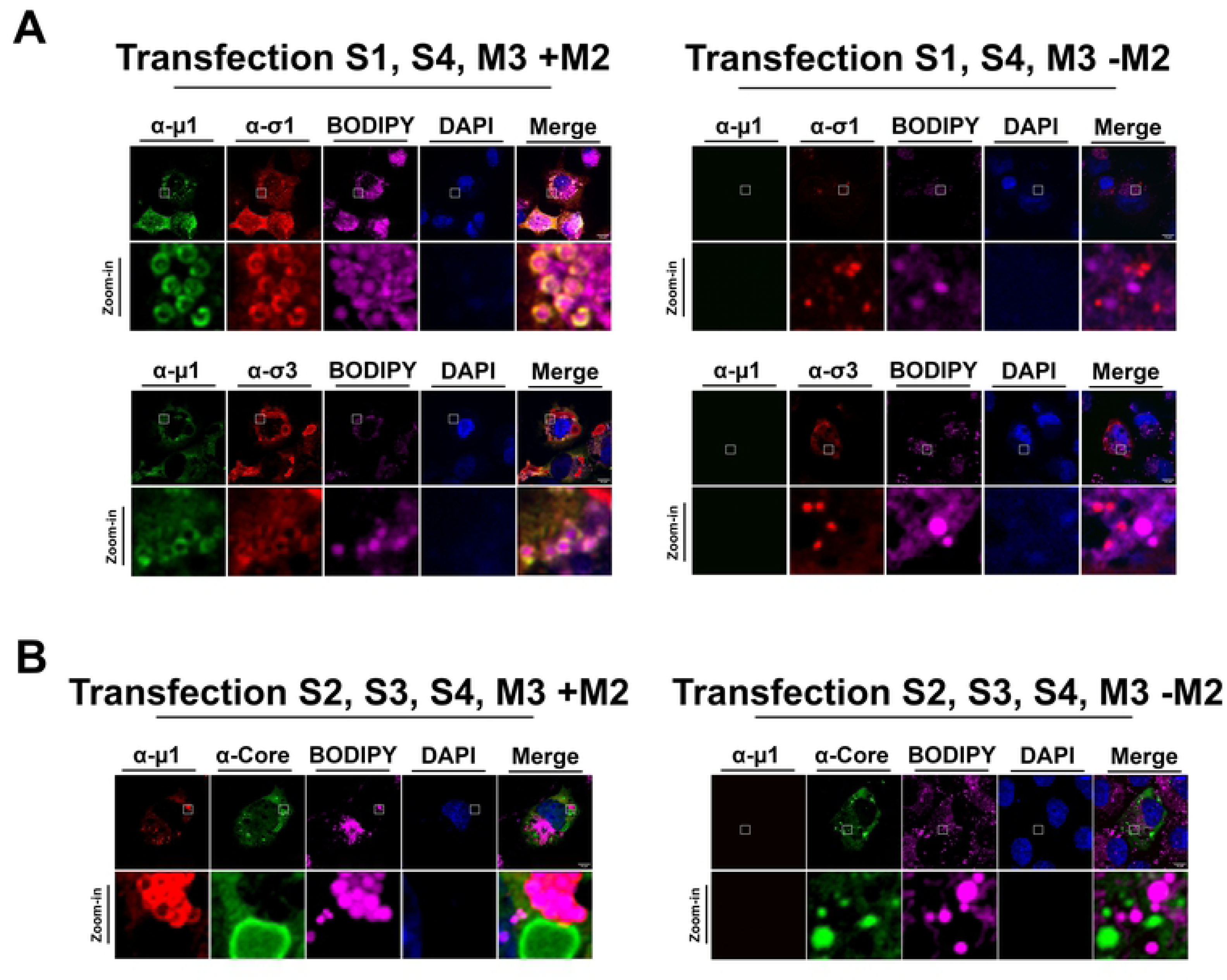
Reovirus outercapsid proteins localize to lipid droplets and perinuclear regions. **(A)** H1299 cells were transfected with S1pcDNA3 (σ1), S4pcDNA3 (σ3), and M3pcDNA3 (μNS) with (left) or without M2pcDNA3 (μ1) (right). Immunofluorescence staining was conducted with antibodies specific to OC proteins μ1 (monoclonal 10F6) and σ1 (monoclonal G5 directly labelled with AlexaFluor 647) (top) or μ1 (10F6) and σ3 (monoclonal 10C1 directly labelled with AlexaFluor 647) (bottom), BODIPY for LDs, and DAPI staining for nuclei. μ1 was detected with secondary antibodies conjugated to AlexaFluor 488. **(B)** H1299 cells were transfected with S2pcDNA3 (σ2), S3pcDNA3 (σNS), S4pcDNA3 (σ3), and M3pcDNA3 (μNS) with (left) or without M2pcDNA3 (μ1) (right). Immunofluorescence staining was conducted with antibodies specific to OC protein μ1 (monoclonal 10F6) and polyclonal rabbit antibodies raised against reovirus cores (α-Core), BODIPY for LDs, and DAPI staining for nuclei. μ1 was detected with secondary antibodies conjugated to AlexaFluor 647, and core protein σ2 with secondary antibodies conjugates to AlexaFluor 488. Represented images were created from Z-stacks acquired using immunofluorescent spinning disk confocal microscopy. Images are representative of at least five images captured for each condition from three biologically independent experiments.

To determine if core proteins localize to LDs in a similar manner to OC proteins, H1299 cells were transfected with the non-structural factory proteins μNS and σNS, core protein σ2, and OC protein σ3 in the presence or absence of μ1. As previously mentioned, μNS is required for pseudo-factory formation but our lab only has access to rabbit polyclonal antibodies for recognizing both μNS and core proteins, rendering discrimination between the two during co-immunofluorescence impossible. However, together with μNS, σNS also forms pseudo-factories for which we have mouse monoclonal antibodies, which allowed us to image both σNS-containing pseudo-factories and core protein σ2 localization. In addition to viral protein staining, BODIPY was used to visualize LDs (Figure 4B). In contrast to the μ1-dependent localization of OC proteins at LDs, core protein σ2 demonstrated no localization at LDs, regardless of μ1 presence.

Altogether, transfection studies revealed that all three OC proteins become LD-associated in a μ1-dependent manner and segregated from core and non-structural proteins. In other words, there is an inherent segregation between OC and core proteins even in absence of virus replication. OC σ3 was previously found to pre-associate with μ1^37, 38^, and our evidence suggests that these proteins likely already pre-associate at LDs. In the fully assembled virion, σ1 is anchored within the cavity of λ2 turrets at each vertex, and therefore it was unexpected that σ1 would also localize to LDs in a μ1-dependent manner. These findings suggest that σ1 may also associate with μ1 or σ3 in infected cells.

### Peripheral factories contain transcriptionally active cores

Having established that OC proteins are spatially segregated from core-only foci, it became important to determine if the core-only regions represented bonafide factories of core amplification versus sites of core protein accumulation without active replication. First, scanning electron microscopy (SEM) array tomography (AT) sample preparation of cell monolayers was conducted using osmium tetroxide as a contrast agent to clearly distinguish lipids and therefore LDs. The SEM AT sample preparation photomicrographs consistently displayed localized areas of virus particle-like regions on both perinuclear-facing and peripheral facing sides of LDs (Figure 5A). Under higher magnification, the perinuclear regions were clearly distinguished as crystalline arrays of fully-assembled viruses. On the peripheral face of LDs, highly contrasted spheres that might represent cores were visible. The SEM AT sample preparation suggested that perinuclear regions represent OC assembled virions, while peripheral foci were likely cores; however, a different EM approach was necessary to achieve certainty that core-like particles were in fact cores. Accordingly, transmission electron microscopy (TEM) using uranyl acetate and lead citrate as a contrast agent was conducted (Figure 5B). While LDs and membranous compartments were not visible in TEM, the proteinaceous content of peripheral versus perinuclear foci was resolved. High magnification photomicrographs were obtained for perinuclear foci (Figure 5B, regions 1 and 2) and peripheral regions (Figure 5B, regions 3 and 4). Visually, particles resembling cores were at the peripheral regions while those resembling whole viruses were perinuclear. To obtain a quantitative conclusion, particle surface area was measured using both ImageQuant image analysis software (Figure 5B, right) and manual diameter measurements (data not shown). Reovirus whole particles are 85nm, while cores are 48nm, and therefore fully-assembled particles are 1.7 times larger than cores. The perinuclear particles, on average, were 1.7 times larger than the peripheral particles, suggesting their identity as whole virions versus cores, respectively. Altogether, electron microscopy indicated that the peripheral regions not only represent areas of core proteins by immunofluorescence, but also contain assembled cores. The perinuclear regions enriched with core and OC proteins by immunofluorescence are enriched in fully-assembled viruses. LDs, which contain OC proteins µ1 and σ3, reside between the peripheral core-containing and perinuclear fully-assembled virus-containing regions.

**Figure 5.**
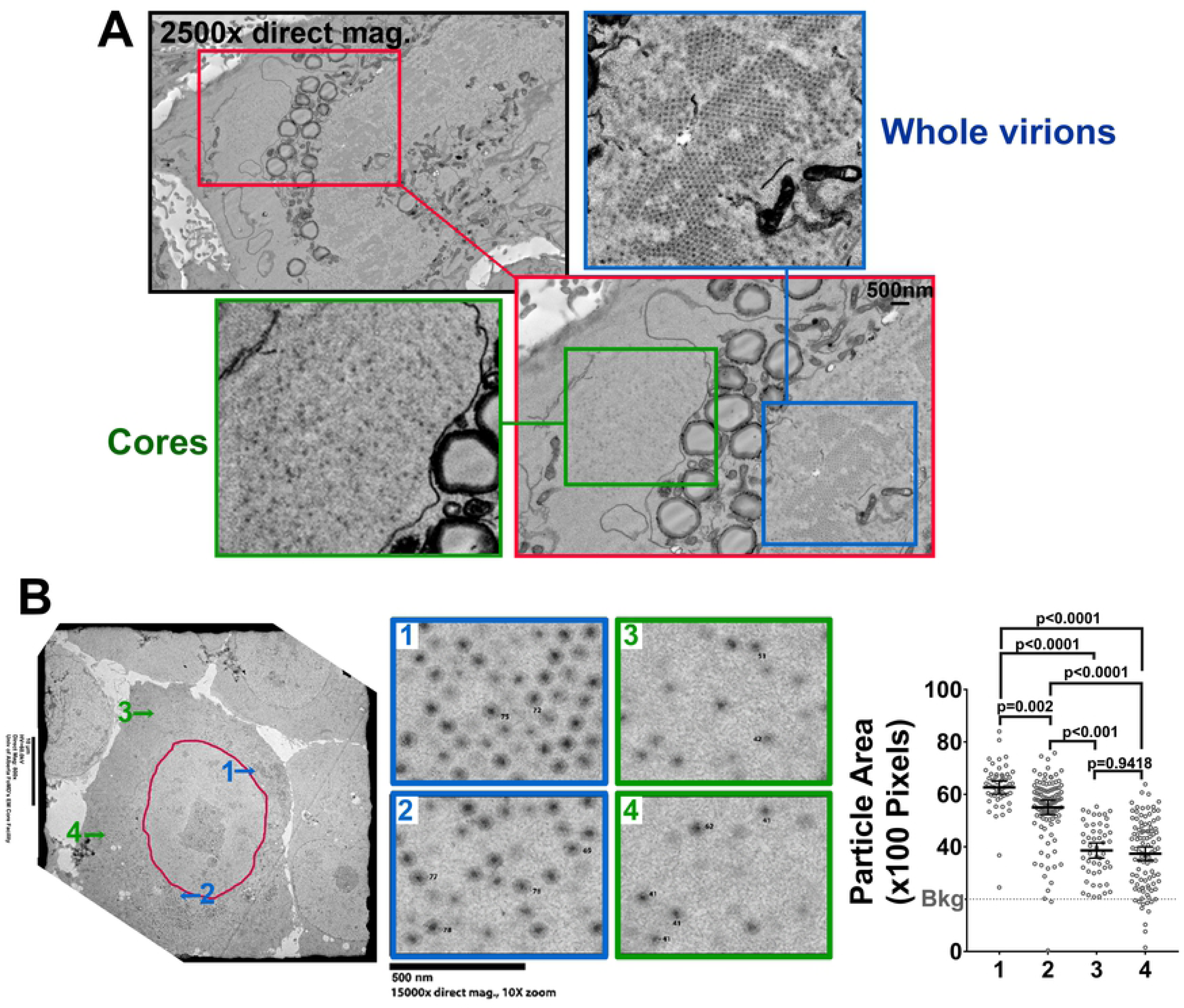
Core-only factories are spatially segregated from whole virion-containing factories. **(A)** Representative SEM image using SEM array tomography sample preparation of an H1299 cell infected with reovirus at an MOI of 3 fixed at 17 hpi. (Green closeup) Distinct region containing cores only. (Blue closeup) Distinct region containing whole viruses. Representative of 8 cells imaged and two independent experiments. **(B)** H1299 cells were infected with reovirus at an MOI of 3, fixed at 16 hpi and imaged via TEM. Images from various regions around the cell were captured at high magnification and the surface area of viral particles in regions 1-4 were measured using ImageQuant image analysis software. Each point represents a single particle (n=239). Statistical analysis is reported as a two-tailed student t-test. Data is plotted as mean +/- 95% CI. **** p<0.0001, *** p<0.001, **p<0.05, ns > 0.05. Representative of 18 cells imaged and 4 independent experiments.

Ultimately, we wanted to know if we could accurately label the core-only regions as “factories”. While the definition of virus factories can vary semantically, herein we define factories as sites of virus replication and/or assembly. Accordingly, to be referred to as factories, core only regions should exhibit *de novo* RNA synthesis and/or active packaging and assembly. In-cell EZ click RNA labelling was applied to monitor *de novo* RNA synthesis in reovirus-infected cells. To focus on viral RNA over cellular RNA, cells had to be treated with actinomycin D (AD) to inhibit host transcription. However, since reovirus replication is also inhibited by AD^39–41^, albeit to lesser extent at low AD concentrations, we needed to wait until reovirus infection was well-established. Cells were therefore co-treated with AD and 5-ethynyl uridine at 15 hpi, fixed at 18 hpi, and stained for core proteins, OC σ1, and *de novo* RNA using click chemistry (Figure 6, Supplementary Figure S6A). Regions positive for core staining but negative of OC staining were consistently positive for *de novo* RNA (Figure 6, Supplementary Figure S6B), indicating that core-only foci are actively synthesizing RNAs and are therefore bonafide factories. The results were striking given that at these late timepoints, packaging of +RNA exceeded *de-novo* synthesis (Figure 2A), and suggested that even late during infection the remaining core-only foci are transcriptionally active. The core+OC-positive perinuclear regions were also positive for *de novo* RNA; but since cells had to be labelled with 5-ethynyl uridine for a minimum of 3 hours for sufficient sensitivity, it remains unclear if *de novo* RNAs were directly produced in perinuclear factories, or whether *de novo* RNA-containing peripheral factories had since travelled to the perinuclear regions. Altogether, data in Figures 5 and 6 indicate that core-only regions are bonafide factories that contain core proteins, cores, and *de novo* RNA synthesis.

**Figure 6.**
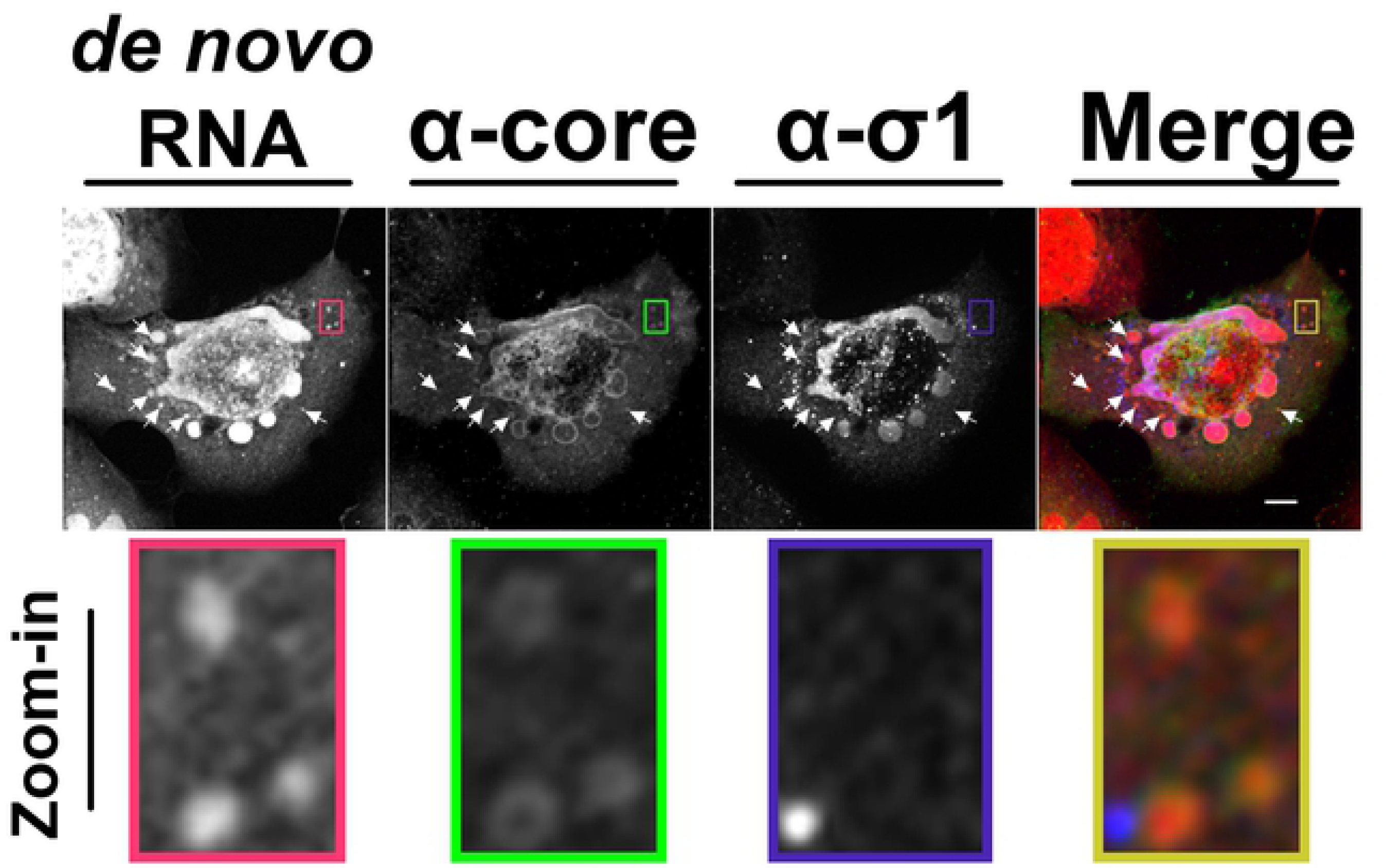
Core-containing factories are transcriptionally active. H1299 cells were infected with reovirus at an MOI of 3 and at 14 hpi, cells were treated with actinomycin D to reduce cell transcription. Between 15 hpi and 18 hpi, cells were stained for *de novo* transcribed RNA using an EZ-click RNA Labelling kit (RNA, red in merged image). Fixed cells were processed for immunofluorescence with rabbit polyclonal α-core antibodies (Alexa Fluor 488, green in merged image) and monoclonal mouse OC protein σ1 antibody G5 (Alexa Fluor 405, blue in merged images). Images were captured by spinning disk confocal microscopy. Scale bar represents 20μm. Highlighted boxes and corresponding close-up images represent example regions positive for core and RNA staining, but negative for σ1. Representative of two independent experiments.

### Temporal and spatial changes to reovirus compartmentalization occur over the course of infection

LD-associated OC proteins, core-only factories and core-and-OC factories were spatially segregated into distinct compartments at 12-18 hpi (Figures 3-6). Given both core and OC RNAs and proteins were synthesized throughout infection (Figure 2), but OC assembly was temporally delayed, we wondered how the compartments progressed over the course of reovirus infection.

Immunofluorescence microscopy was conducted for core and OC proteins over the course of infection in H1299 cells infected at an MOI of 3 (Figure 7 and Supplemental Figure S7). With respect to core-only versus σ3+core-positive factories (Figure 7A and Supplemental Figure S7A), there was progressive transition from exclusively core-only factories at 8 hpi, to both core-only and σ3+core-positive factories starting at 12 hpi onwards. Quantitative analysis using the Volocity 3D image analysis software confirmed that at 8 hpi, despite clear expression of OC proteins (Figure 2), 100% of factories were core-only (Figure 7B and 7C). Over the course of infection, the number of core-only factories decreased over time as a proportion of total factories (Figure 7B) and especially as a proportion of the total overall volume of all factories (Figure 7C). The volume of core-only factories remained similar over time; with an overall average of 1.3µm^3^. Distance of core-only factories from the nucleus decreased from an average of 5.0µm at 8 hpi to an average of 2.0µm at both 12 hpi and 16 hpi, suggesting that core-only factories move towards the nucleus over time (Figure 7D and 7E, respectively). Conversely, shared core+OC factories appeared at 12 hpi, and although the number of core+OC factories remained constant between 12-16hpi (Figure 7B and 7C), core+OC factories grew larger in volume from an average of 4.2µm^3^ at 12 hpi to 19.0µm^3^ at 16 hpi (Figure 7D). Shared core+OC factories were also closer to the nucleus relative to core-only factories at all timepoints, with an average distance of 0.6µm at 12 hpi and 0.2µm at 16 hpi (Figure 7E). Similar characteristics were found when assessing core-only versus σ1+core factories (Supplemental Figure S7B). These results suggest that core-only factories first form in the periphery of the cells but as infection progresses, larger shared core + OC perinuclear factories form where the majority of the viral volume accumulates.

**Figure 7.**
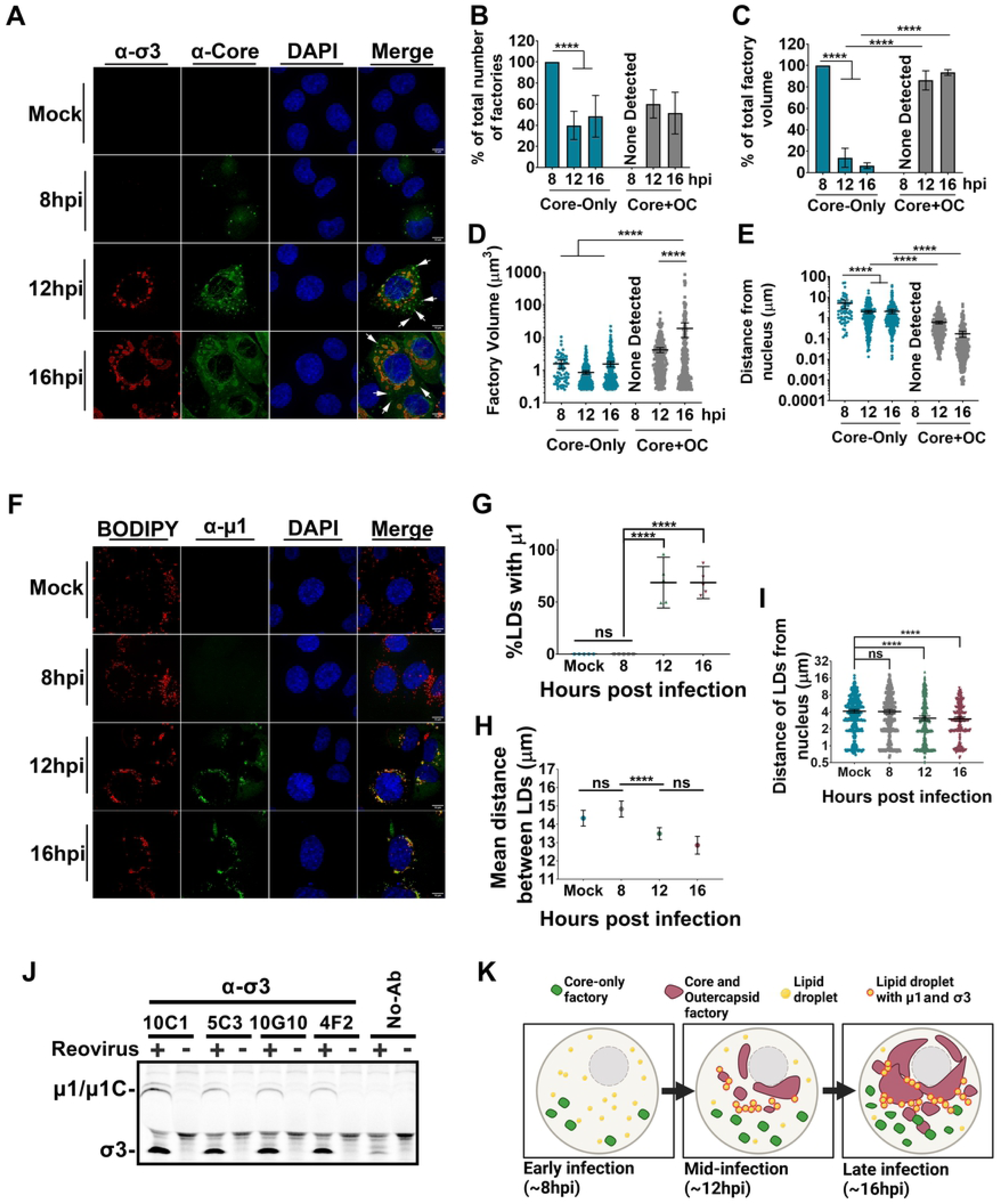
Temporal and spatial changes to reovirus compartmentalization occur over the course of infection. **(A)** Representative images of H1299 cells that were infected with reovirus at an MOI of 3 and fixed at the indicated timepoints. Immunofluorescence spinning disk confocal microscopy was used to capture images of cells stained with monoclonal mouse α-σ3 (10G10, Alexa Fluor 647, red) and polyclonal rabbit α-core (Alexa Fluor 488, green). Cell nuclei were stained with DAPI (blue). White arrows indicate example regions of core-only staining. Images are representative of at least five images for each time point from 3 biological experiments. **(B-E)** σ3/core and σ1/core data were pooled together. At 8hpi there were no visible core- and OC (shared) factories hence labelled “none detected”. **(B)** The number of core-alone or core- and OC (shared) viral objects were quantified and graphed as a percentage of total viral objects within the cell at each timepoint. Statistical analysis is reported as a two-way ANOVA with multiple comparisons between the mean of each column. **(C)** The volume of core-alone or shared factory objects were independently added together for each time point and graphed as a percentage of total viral volume within the cell. Statistical analysis is reported as a two-way ANOVA with multiple comparisons between the mean of each column. **(D)** The factory volume for core-only objects or shared factory objects was found and plotted as individual points at each timepoint. **(E)** The edge-to-edge distance for pixels in each factory type was calculated and plotted at each time point by object identity. **(F)** Representative images of H1299 cells infected with reovirus at an MOI of 3 and fixed at the indicated timepoints. Immunofluorescence confocal microscopy was used to capture cells stained with monoclonal mouse α-μ1 (10F6, Alexa Fluor 647, green). Cell LDs were stained with BODIPY 493/503 dye (red) and nuclei were stained with DAPI (blue). Images representative of at least 5 images for each time point and three biological experiments. **(G)** The percent of LDs within the cell that co-stain with μ1 over the time course. Statistical analysis is reported as aone-way ANOVA with multiple comparisons, comparing the mean of the 8 hpi group to all the others. **(H)** The mean distance between cellular LDs at each time point. **(I)** The edge-to-edge distance of LDs from the nucleus. Statistical analysis was done by one-way ANOVA with multiple comparisons, comparing the mean of the mock group to all the others. Each point represents an individual value from n=5-6 images for each timepoint. **(J)** H1299 cells were infected with T3D^PL^ at an MOI of 3 for 6 hours in the presence of ^35^S-methionine/cysteine. Cell lysates were immunoprecipitated using 5μL of the indicated monoclonal antibodies (σ3: 10C1, 5C3, 10G10, 4F2.) or no-antibody control (No-Ab) prior to resolving by SDS-PAGE. **(K)** Summary diagram depicting factory/protein and LD localization during reovirus infection at ∼8, 12, and 16 hpi. Figure created using Biorender.com. All graphs are plotted as mean +/- 95% CI. Statistical analysis is reported as one-way ANOVA with multiple comparisons between the mean of each column, unless otherwise indicated. **** p<0.0001, *** p<0.001, **p<0.05, ns > 0.05.

With respect to the temporal distribution of OC µ1, the association of µ1with LDs became strongly apparent by 12 hpi onwards (Figure 7F and Supplementary Figure S7C). Approximately 50% of BODIPY-positive LDs were positive for µ1 (Figure 7G), suggesting that µ1 is widely dispersed among LDs of a cell. Several other virus families that utilize LDs for replication and assembly are also described to promote morphological changes to LDs, such as LD aggregation^42^. Similarly, µ1 association altered LD phenotypes; for example, relative to mock-infected cells and cells infected for 8 hours, the cells infected for 12 hpi and 16 hpi exhibited shorter distances between LDs and closer proximities of LDs to the nucleus (Figure 7H and 7I).

It was intriguing that despite similar protein expression kinetics of core and OC proteins at early timepoints (Figure 2), there was minimal detection of OC proteins by immunofluorescence at 8 hpi (Figure 7). Since µ1 and σ3 are known to pre-associate before assembly^37, 38^, co-immunoprecipitation was applied to assess if µ1-σ3 interactions were occurring at early timepoints. Specifically, H1299 cells were infected with reovirus at an MOI of 3 in the presence of ^35^S-methionine/cysteine, and at 6 hpi lysates were subjected to immunoprecipitation with σ3-specific antibodies or a no-antibody control. The same σ3-specific antibodies used for immunofluorescence were herein applied to the co-immunoprecipitations so that similar populations of native proteins could be assessed. As early as 6 hpi, µ1 co-immunoprecipitation with σ3 was observed (Figure 7J).

Moreover, cleavage of µ1 to µ1C occurred, which happens during µ1-σ3 interactions^37, 38^. Accordingly, despite that µ1 and σ3 cannot be detected by immunofluorescence at 8 hpi, these OC proteins were already interacting and were recognized by the same σ3 antibodies used for immunofluorescence analysis of protein native state by 6 hpi. Given the inherent propensity of OC proteins to localize to LDs (Figure 4), we predict that at early timepoints (<12 hpi), OC proteins are not yet sufficiently condensed into specific areas to be detected by immunofluorescence; for example, if instead of being concentrated in a few condensed LDs, OC proteins are dispersed among many LDs and within the cytoplasm, then the dispersed OC proteins at early time points might fail to yield detectable signal by immunofluorescence microscopy.

Altogether, the time course analysis suggested both temporal and spatial regulation of core versus OC localization during reovirus infection (Figure 7K). Specifically, at 8 hpi, despite equivalent expression of core and OC proteins, only core proteins are enriched in peripheral core-only factories where amplification of transcriptionally-active cores occur. Between 6 hpi -12 hpi, OC proteins µ1 and σ3 already pre-associate with each other and at ≥12 hpi, µ1 and σ3 are distributed between LDs and core+OC perinuclear factories where fully-assembled virions reside.

### Peripheral core-only factories transition into the perinuclear regions once established

The time-course analysis revealed spatiotemporal changes to reovirus protein subcellular localization at regular timepoints during infection, but in absence of a live microscopy approach to directly monitor factory movement in real-time, it was difficult to conclude whether (or not) core-only factories transition into perinuclear fully-assembled factories. For example, it was possible that the core-only factories migrated closer to the nucleus over time and fused together until they reached LD-detained OC proteins and transitioned to the perinuclear factories as assembly completes. But it was also possible that core-only factories remain in the periphery and that new cores assemble in new factories closer and closer to the nucleus with increasing propensity to assemble into infectious virions by proximity to LDs where OC proteins reside.

To determine whether peripheral factories remain static or move into perinuclear region as infection progresses, factories were permitted to establish for 10 hpi before cell fixation, or instead, the cells were further incubated for an additional 6 hours with cycloheximide to inhibit *de-novo* virus protein translation and core assembly prior to fixation. The rationale was that cycloheximide treatment would stop the constant flux of core amplification that replenishes peripheral factories, and permit us to capture snapshots of where factories began (10hpi) versus end up (16hpi). If peripheral factories transition towards the nucleus, then there should be many peripheral factories at 10hpi, but depletion of peripheral factories during cycloheximide treatment for the 6 hours, with increased perinuclear factory size. Immunofluorescence confocal microscopy was conducted with α-σNS antibodies to visualize factories independent of core vs OC content (Figure 8A, Supplemental Figure S8A), and with both σNS and OC σ3 to identify factories devoid of OC (Figure 8B, Supplemental Figure S8B). In either case, cells fixed at 10hpi or incubated until 16hpi with DMSO contained an abundance of peripheral factories. But when cells were instead incubated with cycloheximide between 10hpi and 16hpi, there was a loss of peripheral factories in exchange for a predominance of perinuclear factories Altogether, the results suggest that once peripheral-OC-negative factories are established, they then move and merge into larger perinuclear factories (Figure 8C). Moreover, as the peripheral factories move into the perinuclear space, new peripheral factories are established.

**Figure 8.**
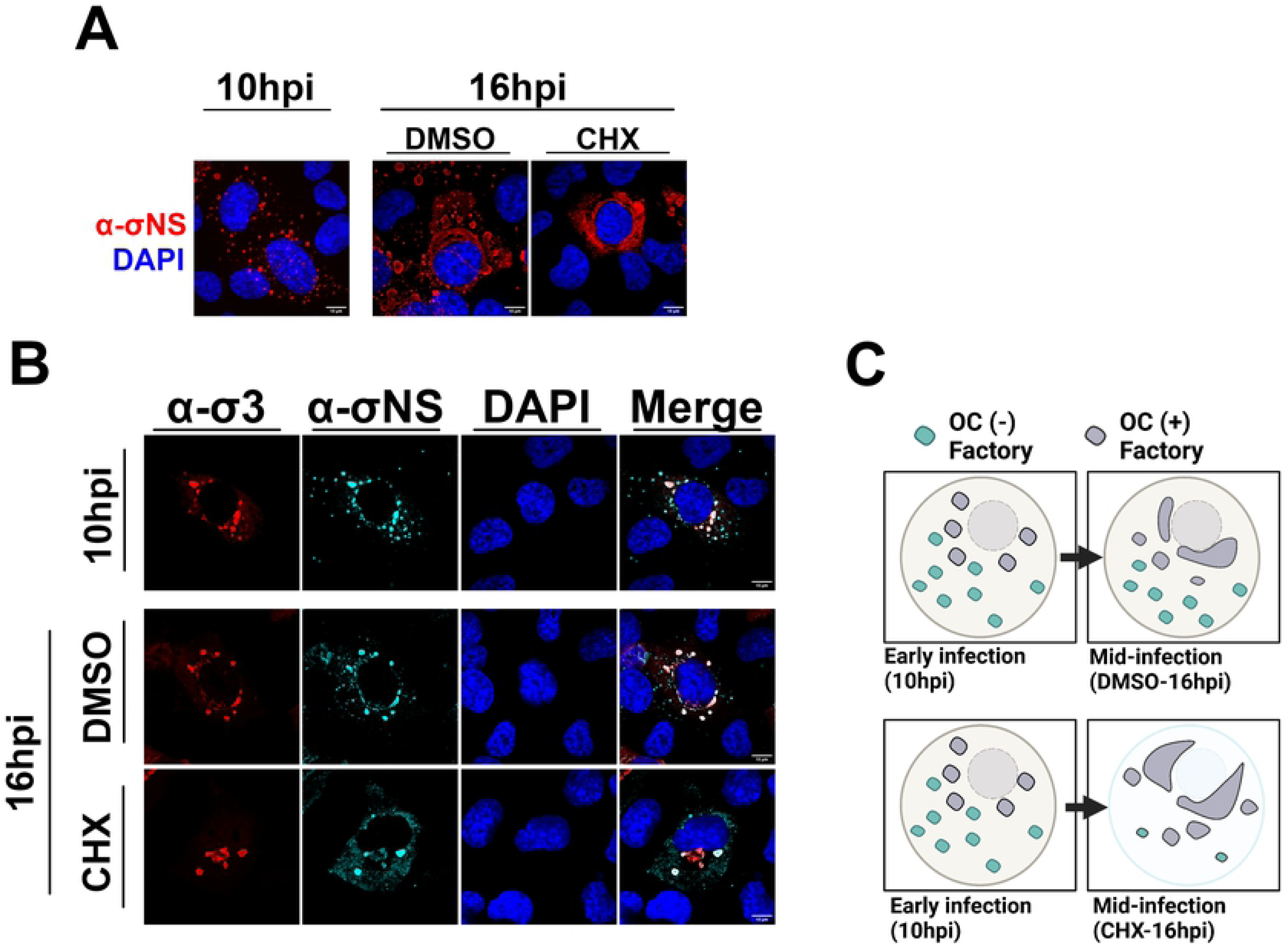
Peripheral viral factories are diminished and diffuse if translation is inhibited during early infection. **(A, B)** Representative images of H1299 cells were infected with reovirus at an MOI of 3, and at 10 hours post infection (hpi) cells were either fixed **(A)** or treated with 100μg/mL cycloheximide or DMSO **(B)**. **(A)** Cells were immunofluorescently labelled with monoclonal mouse α-σNS (3E10 directly conjugated to AlexaFluor 568, red) in combination with DAPI for nuclei staining (blue). **(B)** At 16hpi, cells were fixed and immunofluorescently labelled with monoclonal mouse α-σ3 (10G10, AlexaFluor 647, red), monoclonal mouse α-σNS (3E10 directly conjugated to AlexaFluor 568, cyan) and DAPI for nuclei staining (blue). All images were acquired via immunofluorescence spinning disk confocal microscopy.

### Outercapsid protein μ1 promotes the convergence of peripheral core-only factories into perinuclear core-plus-outercapsid factories

The time course analysis of RNA, protein and virus titers (Figures 1 and 2) versus core and OC intracellular localization (Figure 7) suggested a new working model for temporospatial compartmentalization to orchestrate core versus OC assembly. The next objective was to empirically test the model, and specifically address the role of µ1 and LDs in compartmentalization and assembly. The µ1 protein not only segregates into LDs, but is also the intermediate protein that bridges core particles to the outermost protein σ3. We hypothesized that if µ1 is necessary for distinguished peripheral versus perinuclear factories, then compartmentalization will change in the absence of µ1.

To address the role of µ1 in compartmentalization, the M2 gene that encodes µ1 was silenced with Dicer-substrate short interfering RNAs (DsiRNAs) prior to infection with reovirus (Figure 9 and Supplemental Figure S9). Silencing of M2 did not affect expression of other reovirus proteins, which is to be expected since core amplification can resume without OC proteins (Figure 9A). Silencing of M2 decreased virus titers (Figure 9B), which is also expected because OC proteins are essential for producing the infectious fully-assembled virions. As a negative control, an irrelevant (IRR) DsiRNA was used; this did not affect virus protein expression or titers. As a positive control, siRNAs towards the σ2 core protein-encoding S2 gene were used. Silencing of S2 lead to decreased expression of all viral proteins and reduced titers, as would be expected since the absence of a core protein would diminish core amplification and therefore all stages of reovirus replication (Figures 9A and 9B).

**Figure 9.**
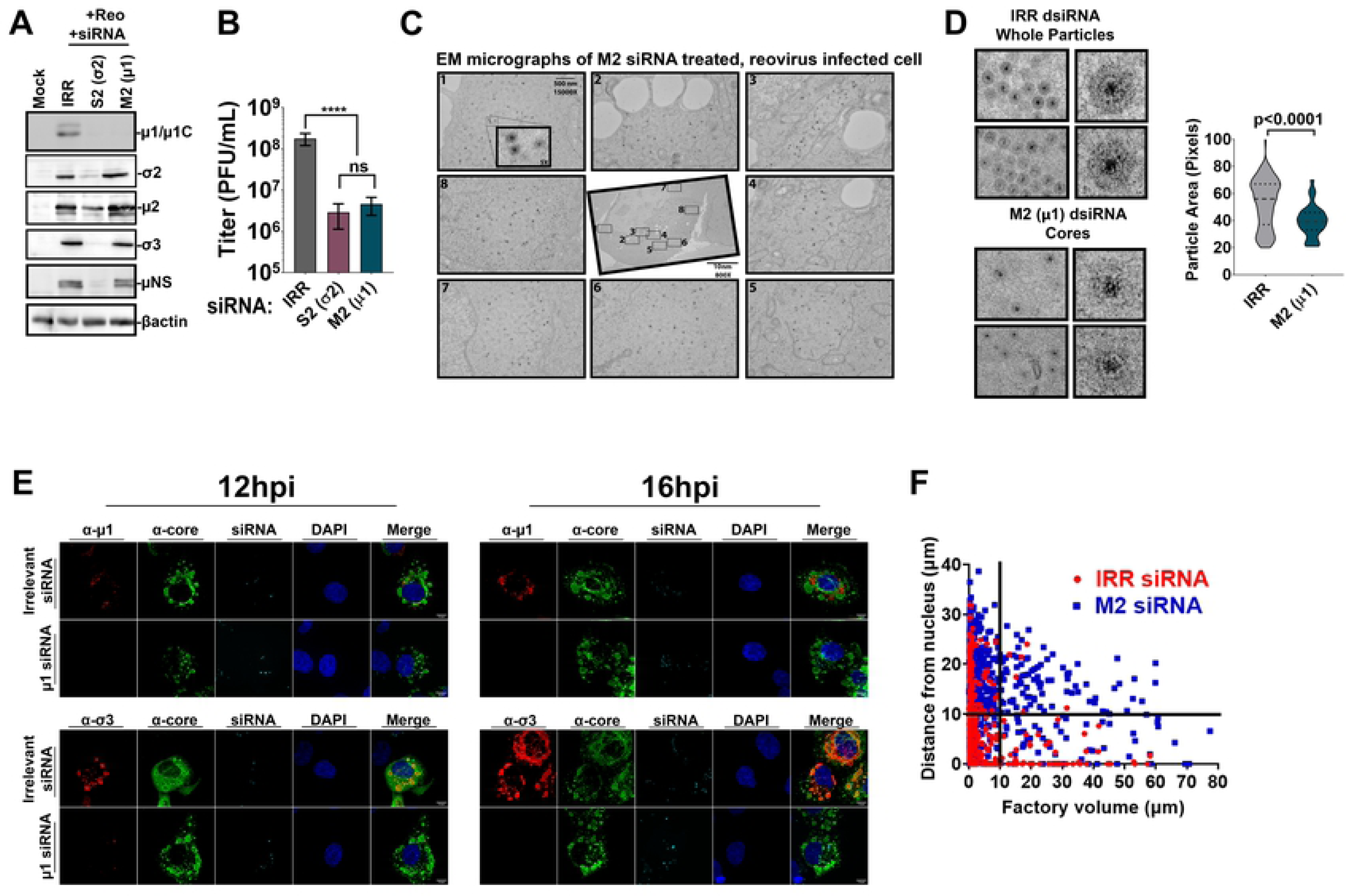
Outercapsid protein μ1 promotes convergence of peripheral core-only factories into perinuclear core-plus-outercapsid factories. **(A-F)** Prior to infecting H1299 cells with reovirus at an MOI of 3, cells were transfected with DsiRNAs: an irrelevant control DsiRNA (IRR), a positive control DsiRNA towards an essential core protein (S2 gene; σ2 reovirus core protein) or test condition (M2 gene; μ1 reovirus outercapsid protein). **(A)** Cell lysates were subjected to SDS-PAGE followed by Western blot analysis to evaluate the extent of gene silencing. **(B)** Lysates from cells infected and pre-treated with DsiRNAs were collected at 18 hpi and viral titers for each condition were assessed by plaque assay. Statistical analysis is reported as a one-way ANOVA with multiple comparisons between the mean of each column. **** p<0.0001, *** p<0.001, **p<0.05, ns > 0.05. **(C)** Cells were fixed at 17 hpi and imaged by TEM. Example images from various regions around a representative M2/μ1 (bottom) DsiRNA-transfected cell. (**D)** Close up of dominant particles found in IRR-versus M2/μ1 DsiRNA-treated cells. Violin plot shows size (area) of reovirus particles imaged by TEM of IRR DsiRNA versus M2/μ1 DsiRNA transfected cells. Statistical analysis is reported as a two-tailed student t-test and graph is plotted as mean +/- 95% CI. **** p<0.0001, *** p<0.001, **p<0.05, ns > 0.05. **(E)** Representative images of DsiRNA-treated infected cells at 12hpi and 16hpi. Cells were immunofluorescently labelled with monoclonal mouse α-μ1 (10F6, Alexa Fluor 647, red) or monoclonal mouse α-σ3 (10G10, Alexa Fluor 647, red) in combination with polyclonal rabbit α-core (Alexa Fluor 488, green), Tye563 for DsiRNA staining (cyan), and DAPI for nuclei staining (blue). **(C-E)** Data from μ1/core and σ3/core co-staining were pooled. **(F)** The volume and distance from the nucleus was measured for each factory in IRR (blue squares) versus M2/μ1 (red circles) siRNA-treated cells, using the α-core channel to capture both core-only and core+OC shared factories. Quadrants were established based arbitrarily at 10µm^3^ volume and 10µm distance from the nucleus to compare ratio of factories between IRR- and M2/μ1 siRNA-treated cells. Data represents eight images per condition and is representative of two independent experiments.

First, TEM of IRR versus M2 DsiRNA treated cells was conducted to establish the effects of M2 silencing on reovirus factories. Similar to normally-infected cells (Figure 5B), IRR DsiRNA transfected and reovirus infected cells exhibited both peripheral core-only and perinuclear core+whole virus factories (Supplementary Figure 9A, IRR siRNA). In contrast, in M2 DsiRNA transfected and reovirus infected cells, there was a predominance of core-containing factories scattered across the cytoplasm that were reminiscent of the core-only peripheral factories in Figure 5B (Figure 9C). Large perinuclear whole-virus containing factories were undetected in M2 DsiRNA treatment. High magnification TEM images confirmed a predominance of fully assembled reovirus particles in IRR DsiRNA treated cells (Figure 9D, top) versus mostly core particles in M2 DsiRNA-treated cells (Figure 9D, bottom), with the expected ∼1.7-fold size differences between cores and whole viruses (Figure 9D, right). The TEM results suggest that when M2 is silenced, not only is there a predominance of cores rather than fully assembled viruses (which is to be expected since OC assembly requires µ1), but importantly that factories now morphologically resemble the core-only factories of standard infections.

Immunofluorescence microscopy at 12 hpi and 16 hpi was then applied to establish the effects of M2 silencing on core and OC protein localization and factory morphology (Figure 9E and Supplementary Figure S10). Tye563-labelled fluorescent siRNA control was used to identify cells that were successfully transfected with DsiRNAs. As expected, in M2 silenced cells, µ1 staining was too low to detect. The reovirus σ3 was still excluded from core factories, and was either in small, segregated foci or below detection, which may suggest that in absence of intermediate µ1 protein to bridge σ3 onto cores or onto LDs, σ3 becomes diffuse and aggregated (Figure 9E and Supplementary Figure S10). Importantly, the phenotype of core-positive factories was changed in M2-silenced cells. Rather than the accumulation of both peripheral and perinuclear factories as seen in IRR treated cells, M2 silencing led to larger core factories that remained further away from the nucleus (Figure 9F and Supplementary Figure S9B). Specifically, while 55% of factories were within 10µm distance from the nucleus in IRR DsiRNA-treated cells, only 26% of factories were within that proximity when M2 was silenced. With respect to volume, both M2 and IRR silenced cells had similar numbers of factories >10µm^3^ (22% for M2 and 17% for IRR). Altogether, results from the TEM and immunofluorescence analysis suggest that µ1 is the linchpin not only for OC assembly, but for the convergence of peripheral core-only factories into perinuclear core+OC factories.

## DISCUSSION

For *Reoviridae* members that have both a core and OC, the premature assembly of the OC would halt amplification by transcriptionally-active progeny cores. Accordingly, we hypothesized that core amplification and OC assembly were orchestrated. Indeed, kinetic analysis of *de novo* RNA synthesis (core amplification) versus infectious titers (OC assembly) for reovirus indicated an ∼3hr delay between these processes (Figure 1). There were no delays in the synthesis of mRNAs encoding OC proteins, nor in the expression of the OC proteins (Figure 2), indicating that delayed OC assembly was not a consequence of delayed availability of OC proteins. Instead, we discovered that early during infection, when core amplification presided, there were core-only factories containing bonafide core particles and *de novo* transcription (Figures 3-7). Meanwhile OC proteins accumulated at LDs (Figure 4). As infection progressed to timepoints where infectious virions were detected, factories containing core and OC proteins developed proximal to the nucleus. During cycloheximide treatment, peripheral core-only factories turned into nuclear factories (Figure 8), implying a continuous renewal of peripheral factories albeit to lower proportions as infection progresses. Moreover, silencing of the OC protein µ1 decreased the formation of perinuclear factories (Figure 9), suggesting the transition to perinuclear factories is promoted by µ1. Altogether this generates a model whereby early during infection, core amplification occurs in peripheral core-only factories while OC proteins traffic to the LDs, which are nearer to the nucleus (Figure 10). As infection progresses, core particles become encapsulated by OC proteins near LDs until fully assembled virions congregate in large perinuclear factories. Beyond 12hpi, the predominance of new virus produced is immediately whole infectious virus, using pools of core-derived +RNAs and continued protein translation from LD-proximal factories.

**Figure 10.**
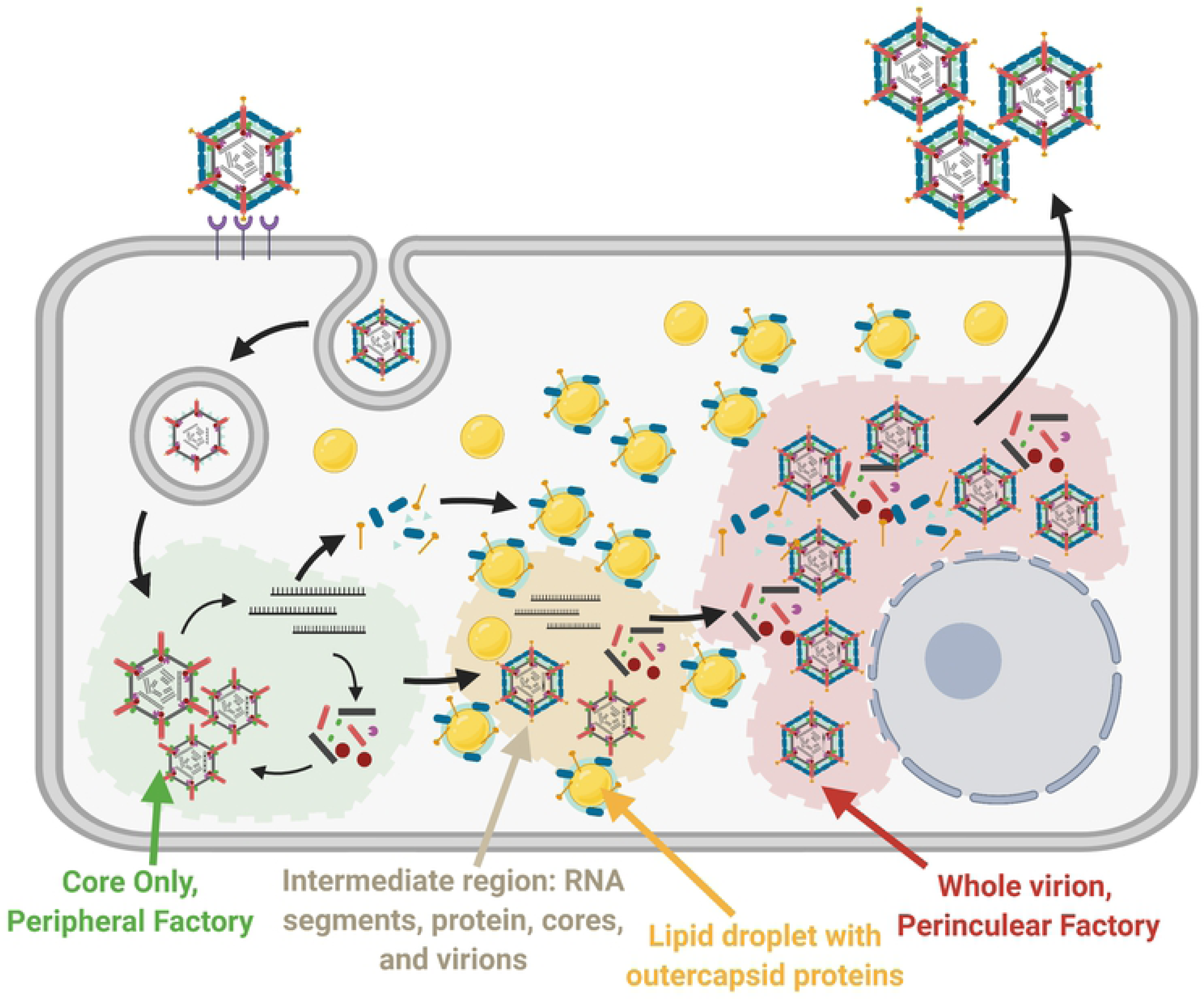
Reovirus assembly is spatially and temporally compartmentalized. A cartoon model summarizing the findings of our study. There exist four segregated areas of reovirus assembly: 1) Core-only peripheral factories, 2) Intermediate regions where residual amplified RNA is translated to proteins and assembled into particles, 3) OC proteins on LDs, and 4) whole virions and core and OC proteins in perinuclear factories. After entry, core particles undergo primary transcription and translation in peripheral core-only factories. Newly synthesized proteins assemble into progeny cores, which then undergo secondary rounds of transcription and translation. As infection progresses, μ1 facilitates OC protein accumulation on LDs, and perinuclear factories form containing predominantly assembled whole virions. Figure created using Biorender.com

The creation of core-containing peripheral factories is most likely by active inclusion, while the absence of OC proteins in core-only factories is likely by active exclusion. In support of active inclusion of core proteins in core-only factories, several studies have demonstrated that the non-structural protein µNS functions as a scaffold to bind core proteins σ2, µ2, λ1, λ2 and λ3^22–25^. µNS also binds the non-structural protein σNS, which binds RNA. One can therefore imagine these basic components producing active enrichment of all components necessary to generate progeny core particles that amplify replication. Unlike the perinuclear factories that contained tightly aggregated crystalline structures of fully assembled viruses, the core-only factories contained more dispersed core particles; this makes sense given that cores generate *de novo* RNA and are hence unlikely to be tightly arranged.

Our findings support active exclusion of OC proteins from peripheral factories, since no OC proteins were present in core-only factories despite their high expression and known ability to interact with cores. For example, the OC protein σ3 also binds viral dsRNA to inhibit stimulation of host innate defense mechanisms^43–45^, yet σ3 is not found in core-only factories. While µ1 was previously found on LDs and µ1-σ3 are known to interact, this is the first time that all three OC proteins were found to concentrate on LDs in a µ1-dependent manner and away from µNS-dependent neo-organelles. The accumulation of OC proteins in LDs support that these proteins are instead trafficked away from sites of core-only factories. The molecular details of how all three OC proteins converge at LDs might in future reveal the precise mechanism of OC assembly. While we can not ascribe intelligence to viruses, there is an astonishing evolutionary brilliance of using LDs and µ1 as the linchpins to orchestrate core versus OC assembly. The OC µ1 is the intermediate between cores and the OC protein σ3, so withholding µ1 would be a “one stop” mechanism to most-rapidly delay all OC assembly. As µ1 at LDs concentrates the remaining OC proteins σ3 and σ1, this could become a mechanism to make OC assembly more efficient than stochastic assembly of dispersed cytoplasmic proteins. In the future it would be interesting to establish if LDs do in fact function directly as a “one stop” depot for OC protein assembly onto cores, or whether they merely act as a sink to sequester OC proteins and delay full assembly. Moreover, previous studies found that the folding of OC protein σ3 and its association with µ1 uses the TRiC/CTT chaperone network^46^, while folding of OC σ1 requires chaperone Hsp90^47^; it would be fascinating to know if the host chaperones are involved in active exclusion of OC proteins from core-only factories, for example by transporting the OC proteins towards the nucleus.

The core-only factories transitioned towards the nucleus when cycloheximide was employed to stop further virus amplification. Previous studies found that the µ2 protein associates with microtubules^48–50^ and the current dogma is that these interactions mediate movement of viral factories. However past studies focused on late timepoints during infection when perinuclear factories predominant, and it would be interesting to revisit microtubule-µ2 associations with respect to whether they also support the movement of core-only factories. It would also be very interesting to establish if there are differences among distinct strains of reovirus with respect to temporospatial compartmentalization of core and OC proteins. For example, it was previously found that a mutation in the µ2 protein can affect whether factories at late timepoints of infection are globular or filamentous^49, 51, 52^. Specifically, in strains of reovirus such as T3D^PL^ used in our studies, the perinuclear factories are filamentous (large amorphous inclusions and thread-like staining), whereas for a closely related laboratory strain T3D^TD^, perinuclear factories are globular (smaller and rounder separated inclusions without a filamentous appearance). In the future it would be interesting to establish if strains of reovirus differ in the kinetics of core-only factory establishment and movement towards the nucleus to help create the filamentous versus globular morphologies.

Even more intriguing than potential differences among strains of reovirus are the evolutionary similarities versus differences among unique genera of *Reoviridae,* with respect to overcoming the replication-versus-full assembly conundrum. The *Reoviridae* family diverges into two subfamilies: members of the *Spinareovirinae,* such as reovirus, have turret-like structures at each 5-fold axis made from pentameric core λ2 proteins that grasp and project the trimeric OC σ1 cell attachment protein. In contrast, rotavirus, as an example of the non-turreted members of the *Sedoreovirinae* subfamily, has attachment-binding protein VP7 embedded in the OC major protein VP7. Although similar to reovirus µ1, rotavirus has an intermediate OC protein VP6 and encodes 6, rather than 2, non-structural proteins. Similar to the reovirus non-structural proteins µNS and σNS, which are involved in virus factory formation, rotavirus encodes NSP2, 5, and 6 for this same purpose. But in rotavirus, it is the factory-forming NSP2, 5 and 6 that associate with LDs to nucleate virus factories^53–55^; in contrast to reovirus that seems to usurp LDs instead for OC localization. Also, unlike reovirus, rotavirus encodes an ER-directed membrane protein NSP4 and the OC proteins VP4 and VP7. For rotavirus, full virus assembly requires budding into the endoplasmic reticulum and then release via nonclassical vesicular transport. Accordingly for rotavirus, the addition of the OC is clearly spatially regulated at the ER. Although to our knowledge, the precise mechanism of maintaining rotavirus cores “as cores” and to delay OC assembly has yet to be directly addressed, one might imagine that similar to reovirus, there might be core-only factories that are distinct from the core-containing factories at the ER awaiting full assembly. It seems that during evolution, members of *Reoviridae* shared in common the utilization of subcellular structures, such as LDs, to segregate factory-forming, core and OC proteins; but it is fascinating that while rotaviruses evolved to use LDs to organize the factory-forming non-structural proteins^53–55^, reovirus instead evolved to use the same organelle to organize OC proteins. Even more intriguing is that LD association has been reported for non-structural proteins of viruses other than rotavirus, such as those in the *Flaviviridae*^56, 57^ and *Bunyaviridae*^58, 59^ families; why did reovirus evolve to sequester OC proteins instead of non-structural proteins on LDs?

The parting questions are whether temporospatial compartmentalization for reovirus occurs similarly in the different cell types it can infect, whether association of OC proteins with LDs serves to delay OC assembly or whether it conveys other advantages to reoviruses. As for potential differences among host cells, reovirus naturally spreads by the fecal-oral route and infects intestinal enterocytes and M cells. LDs are single membrane vesicles that store neutral triacylglycerol. Enterocytes can be enriched with LDs by absorption from dietary sources or intracellular synthesis, and are involved in the distribution of fatty acids to the rest of the body^60–63^. As such, reovirus would have access to LDs during natural infection. But reovirus is also undergoing clinical testing as a candidate cancer therapeutic since it replicates efficiently in cancer cells. In general, untransformed cells preferentially obtain fatty acids from extracellular sources, while metabolic changes in cancer cells often support enhanced *de novo* fatty acid synthesis and LD formation^64, 65^. While major determinants of enteric and tumor specificity have been described^19–21, 66–68^, could additional host factors, such as LD abundance, also contribute to how well reovirus thrives in these environments? If so, then does diet affect natural infection by reovirus and its pathogenic cousins? Could differences in LD levels affect the potency of reovirus oncolysis? Finally, apart from storage depots for fatty acids, LDs also contribute to maintaining cell health during stress by appropriating harmful lipids, maintaining membrane homeostasis, helping clear unfolded ER proteins that could aggravate unfolded protein responses and sequestering cellular proteins^69^. Accordingly, it is not a surprise that viruses use LDs not only for assembly, but for other advantages. For example, hepatitis C virus exploits the mobile nature of LDs, like ships, to move towards regions dense in virus replication^56^. Reovirus seemed to also cause redistribution of LDs (Figure 7); might this confer transport advantages to reovirus factories? The association of reovirus µ1 on LDs was previously also found to affect cell death^26^; but might there yet be additional roles for associating σ3 and σ1 with LDs?

In conclusion, the original model of reovirus replication showing core amplification and OC assembly occurring in shared factories left a conundrum, since OC assembly would thwart further core amplification. The temporospatial compartmentalization of replicating cores at core-only factories, shuttling of OC proteins to LDs, the complete assembly at perinuclear LD-proximal factories and the associated 3-hour delay between peek core amplification versus full assembly, implores a more complex model of orchestrated reovirus assembly that overcomes the replication-versus-assembly conundrum, as described in our summary model (Figure 10).

## MATERIALS AND METHODS

### Cell lines, viruses, and plaque assay

H1299, T47D, MCF7 and L929 cells were purchased from the American Type Culture Collection (ATCC) and maintained at 37°C with 5% CO_2_. Adherent L929s were cultured in MEM (M4655, Millipore Sigma) supplemented with 5% FBS (F1051, Millipore Sigma), 1× non-essential amino acids (M7145, Millipore Sigma) and 1mM sodium pyruvate (S8636, Millipore Sigma). Remaining cells were cultured in RPMI (R8758, Millipore Sigma) supplemented similarly to L929 (H1299s in 5% FBS and T47D/MCFs in 10% FBS). L929s cultured in suspension were grown in Joklik’s modified MEM (pH 7.4) (M0518, Millipore Sigma) supplemented with 2g/L sodium bicarbonate (BP328, Fisher Scientific), 1.2g/L HEPES (BP310, Fisher Scientific), 1× non-essential amino acids (M7145, Millipore Sigma) and 1mM sodium pyruvate (S8636, Millipore Sigma). Reovirus serotype type 3 Dearing-PL (T3D^PL^; Dr. Patrick Lee, Dalhousie University) was propagated in suspension L929 cells from a seed stock to preserve genetic identity, extracted with Vertrel XF (Dymar Chemicals) three times, and purified by ultracentrifugation on cesium chloride gradients, as previously described^70^.

### Plaque Assays

For reovirus titers, plaque assays were done in L929 cells. Reovirus dilutions in serum-free media were added to confluent monolayers of L929 cells for 1 hour with gentle rocking every 10 minutes. After 1 hour, an agar overlay was added (1:2 ratio of 2X JMEM media, 1:4 ratio of complete MEM media, and 1:4 ratio of 2% agar). Overlays were allowed to solidify at room temperature for 30 minutes before incubation at 37°C/5% CO_2_ for 4 days. Then, 4% paraformaldehyde (PFA; in PBS) was added to the overlay for 1 hour at room temperature. PFA was then discarded and agar overlays were gently removed. Cells were then fixed with methanol for 15 minutes at 4°C.

After discarding the methanol, plaques were visualized by staining with a 1% (wt/vol) crystal violet solution and plaques were manually counted.

### Cloning

Reovirus RNA was extracted from infected L929 cells and was used as a template in cDNA synthesis using gene-specific primers. Reovirus genes were amplified by PCR using gene-specific primers and were cloned into the vector pcDNA3 using the restriction sites KpnI and NotI (S1, S2, S3, S4, M1, M3, L1, L2, L3) or EcoRV and NotI (M2).

### New antibodies

Reovirus cores were generated by treatment of purified viruses with chymotrypsin as previously described^33^, purified by CsCl, dialyzed against 10mM Tris pH 7, 150mM NaCl, and UV-inactivated. Antibodies were generated in rabbits by ProSci Incorporated using our aliquots of 2mg/ml x 700ul x 2vials. Rabbit antibodies to reovirus specific reovirus proteins were generated by biologic Corp from IPTG induced BL21 (DE3) bacterially expressed N-His6 tagged and purified whole proteins.

### Western blot

For standard Western blot analysis, cells were first washed with 1xPBS before lysing with RIPA buffer (50mM Tris pH 7.4, 150mM NaCl, 1% NP-40, 0.5% sodium deoxycholate) supplemented with protease inhibitor cocktail (P8340, Sigma). After collection, 5x protein sample buffer (250mM Tris pH 6.8, 5% SDS, 45% glycerol, 9% β-mercaptoethanol, and 0.01% bromophenol blue) was added to a final concentration of 1x. Samples were then boiled at 100°C for 10 minutes. Samples were electrophoresed on 10% SDS-acrylamide gels. Following SDS-PAGE, proteins were transferred from the gel onto nitrocellulose membranes using a Trans-Blot Turbo Transfer System (Bio-Rad). Membranes were incubated with block buffer (3% newborn calf serum (NCS)/TBS-T) for 1-2 hours at room temperature, followed by an incubation with a primary antibody solution (block buffer containing: rabbit pAB against whole virus at 1:10,000 (Biologics International Corp), rabbit pAB against core at 1:1000 (Biologics International Corp), mouse α-σ3 (4F2) at 1:1000 (Developmental Studies Hybridoma Bank (DSHB)), mouse α-μ1 (10F6) at 1:1000 (DSHB), rabbit α-μNS at 1:1000 (Biologics International Corp) and/or mouse α-σ1 (G5) at 1:500 (DSHB)) for either 1 hour at room temperature or overnight at 4°C followed by a secondary antibody solution (block buffer containing: goat α-rabbit AlexaFluor 488 or 647 at 1:1000, goat α-rabbit HRP 1:10,000, goat α-mouse AlexaFluor 488 or 647 at 1:1000, and/or goat α-mouse HRP 1:10,000) for 1 hour at room temperature. Membranes were washed 3x for 5 minutes each in TBST between primary and secondary antibody incubations. Prior to imaging, membranes with HRP-conjugated secondaries were incubated with ECL Plus Western Blotting Substrate (32132, Thermo Fisher Scientific) for 5 minutes at room temperature. Membrane imaging was carried out using an ImageQuant LAS4010 Imager (GE Healthcare Life Sciences). Densitometric analysis was performed with ImageQuant TL (GE Healthcare Life Sciences) software, and images were processed for display in Adobe Photoshop. All incubation steps for membranes were done with gentle rocking.

### S^35^-pulse labelling, immunoprecipitation, virus purification, and gel analysis

For labelling *de-novo* protein translation, H1299 cells were cultured in the presence of S^35^-methionine and cysteine (10µCi/ml, NEG772007MC, Perkin Elmer) in methionine/cysteine free MEM at indicated time and duration. Cell lysates (4cm^2^) were collected with RIPA buffer (50mM Tris pH 7.4, 150mM NaCl, 1% NP-40, 0.5% sodium deoxycholate) supplemented with protease inhibitor cocktail (P8340, Sigma). Lysates were incubated on ice for 30minutes and cleared of nuclei and debris by centrifugation at 2K RPM for 1 minutes. For immunoprecipitation or co-immunoprecipitation, per sample, 25μL of protein G magnetic beads (LSKMAGG10; Millipore) were used. Beads were washed 2x with Co-IP buffer (50 mM Tris-HCl (pH 7.4), 150 mM NaCl, 0.8% NP-40, with protease inhibitor cocktail added prior to use) prior to incubation with 5μL of primary antibody (σ3: 10C1, 5C3, 10G10, 4F2, or polyclonal anti-reovirus antibodies) for 2 hours at room temperature with gentle agitation. Beads were then washed 3x with co-IP buffer. After washing, the bead-antibody mixture was added to samples in equal amounts and allowed to incubate for 2 hours at room temperature. Beads were then washed 3x with co-IP buffer, and subjected to SDS-PAGE. For purification of radiolabelled viruses, cleared RIPA lysates were overlayed on 500 μL 1.33g/cc CsCl and subjected to 100,000 x g for 2 hours. Pellets were resuspended in protein sample buffer and subjected to SDS-PAGE. Gels were dried to filter paper and exposed to phosphor-screens prior to imaging using a Typhoon™ 5 laser-scanner platform (Cytivia). Quantification was performed by densitometry using with ImageQuant TL (GE Healthcare Life Sciences) software. Images were processed for figures in Adobe Photoshop.

### RT-qPCR

Cells were lysed in TRI Reagent® (T9424, Millipore Sigma) and the aqueous phase was separated following chloroform extraction as per the TRI Reagent® protocol. Isopropanol was mixed with the aqueous phase and RNA was isolated as per the GenElute Mammalian Total RNA Miniprep kit (RTN350, Millipore Sigma) protocol. RNA was eluted using RNAse free water and total RNA was quantified using a NanoDrop spectrophotometer.

Using 1µg RNA per 20µl reaction, cDNA synthesis was performed with random primers (48190011, ThermoFisher Scientific) and M-MLV reverse transcriptase (28025013, ThermoFisher Scientific) as per the manufacturers protocol. Following a 1/4 cDNA dilution, RT-PCR reactions were executed following the Sybr Select (4472920, Invitrogen) protocol using reovirus gene-specific primers (listed in **Supplementary Figure 11**) and the CFX96 system (Bio-Rad). All RT-qPCR reaction plates included a no template and no reverse transcription control.

### Immunofluorescent confocal microscopy

*Sample preparation*. 5x10^4^ to 1x10^5^ H1299 (or T47D or MCF7) cells were plated on 18mm glass coverslips in 12-well plates and allowed to adhere for ∼24 hours. Viruses were diluted in serum-free media and added to wells for 1 hour at 37°C, with gentle rocking every 10 minutes. Virus media was then removed, replaced with complete media, and cells were incubated at 37°C until the required hpi. Media was removed and the cells were washed with PBS prior to fixation with pre-warmed 4% PFA for 30 minutes at room temperature. PFA was discarded and the cells were washed 3x for 5 minutes each with PBS. Cells were then permeabilized by incubation with 0.1% Triton X-100 (Tx100) in PBS for 5 minutes at room temperature. PBS-Tx100 was then discarded and cells were washed 3x for 5 minutes each with PBS. After permeabilization, cells were then incubated with blocking solution (3% NCS/PBS with 0.1M Glycine) for 1 hour at room temperature. Then, blocking solution was removed and cells were incubated with primary antibody solutions (block buffer containing: rabbit pAB against whole virus at 1:1000 (Biologics International Corp), rabbit pAB against core at 1:1000 (Biologics International Corp), mouse α-σ3 (4F2) at 1:250 (DSHB), mouse α-σ3 (10G10) at 1:250 (DSHB), mouse α-μ1 (10F6) at 1:1000 (DSHB), rabbit α-μNS at 1:1000 (Biologics International Corp), mouse α-σNS (2F5) at 1:500 (DSHB), mouse α-σ1 (G5) at 1:250 (DSHB), mouse α-σ3 (10C1 or 10G10) directly labelled with AlexaFluor 647 using the Apex AlexFluor 647 Ab-Labelling Kit (A10475, Invitrogen) at 1:100, mouse α-σ1 (G5) directly labelled with AlexaFluor 647 using the Apex AlexFluor 647 Ab-Labelling Kit, (Invitrogen) at 1:100, or mouse α-σNS (2A9) directly labelled with AlexaFluor 568 using the Apex AlexFluor 568 Ab-Labelling Kit, (A10494, Invitrogen) at 1:100 overnight at 4°C. Directly conjugated antibodies were made according to the manufacturers instructions. Primary antibody solution was then removed, and cells were washed with 0.1% Tween-20 in PBS 3x for 5 minutes each. For secondary antibody staining, antibodies (Alexa Fluor 647 goat α-rabbit, Alexa Fluor 488 goat α-rabbit, Alexa Fluor 647 goat α-mouse, Alexa Fluor 488 goat α-mouse, and Cy3 goat α-mouse) were then diluted (1:300) in blocking solution and cells were allowed to incubate in the solution for 1 hour at room temperature while blocked from light. Cellular dyes, such as BODIPY 558 (5μM), D3835, Invitrogen) or BODIPY 493/503 (5μM, D3922, Invitrogen) and DAPI (0.1μg/mL, D1306, Invitrogen) or Hoechst (1μg/mL, H1399, Molecular Probes) either added in together with the secondary antibody incubation or performed after, following manufacturer instructions. Coverslips were then mounted onto glass slides using 1 drop (∼50μL) of Prolong Diamond AntiFade Mountant (Invitrogen). The edges of the coverslips were then sealed with nail polish. Slides were allowed to dry overnight at room temperature, protected from light, prior to imaging.

#### Image acquisition

All confocal images were captured using the University of Alberta Cell Imaging Core’s spinning disk confocal microscope (Quorum Technologies) with the following lasers: 405nm, 491nm, 561nm, 642nm and corresponding emission filters for DAPI, GFP, TRITC and Cy5. Images were acquired with a 60X/1.42NA oil-immersion lens on a Hammamatsu C9100 EMCCD camera (Hamamatsu Corp) using Volocity (Quorum Technologies) software.

#### Image analysis

For visual presentation, images were processed from Z-stacked images projected into one 2D-image based on maximum intensities in ImageJ (NIH) software with the Fiji plugin. The brightness of each channel was adjusted for display purposes only, and images were prepared for figures in Adobe Photoshop. For quantification, 3D-unedited images were used in Volocity software. To quantify the number of a certain object, as well as the objects volume, Volocitys compartmentalization was used to select objects >0.25μm^3^ in size alongside visually thresholding to ensure only true factory objects were selected, not background signals. To determine the distance from the nucleus of each object, a circle was drawn around the nucleus to create a region of interest (ROI) where the edge-to-edge distance from a particular object to the ROI could then be measured. “Shared” objects (OC with core) were determined to be objects compartmentalized within each other (ie. when a core object was found within an OC object, or vice versa).

### Electron microscopy

Electron microscopy and sample preparation was performed at the University of Alberta’s Cell Imaging Core. In brief, cell monolayer samples were prepared on 13mm diameter Thermanox coverslips and fixed with 2.5% glutaraldehyde in 0.1M cacodylate buffer. Contrasting was performed using osmium tetroxide and post-staining with uranyl acetate and lead citrate. Tissue processing was performed using an automated Leica EM TP Tissue Processor. The Thermanox coverslips were embedded on BEEM capsules and thermally polymerized at 65 °C for 24 hours. Seventy nanometer (nm) thick ultrathin sections were made by a Leica EM UC7 ultramicrotome and diamond knife. Four nm thick carbon evaporation was performed using a Leica EM ACE600 high vacuum coater. The ultrathin sections were imaged under a Hitachi H-7650 Transmission Electron Microscope at 80keV high tension. For SEM array tomography imaging, 50 nm thick serial ultrathin sections were made by a Leica EM UC7 ultramicrotome and diamond jumbo knife and mounted on a Si wafer (10 x 10 mm^2^). Five nm thick carbon evaporation was performed and imaged under a Hitachi S-4800 field emission gun scanning electron microscope at 5keV high tension.

### EZ-Click (+)RNA labelling

*De novo* RNA labelling was performed using the EZClick™ Global RNA Synthesis Assay Red Fluorescence Kit (K718, Biovision) according to manufacturer’s instructions. Four hours prior to fixation, cells were preincubated with the kit supplied actinomycin D to suppress nuclear transcription to allow the visualization of viral RNA synthesis. It should be noted that co-staining using 647 fluorescent secondary antibodies was incompatible in conjunction with the kit, likely due to a quenching effect on the 647 fluorophores due to the labelling procedure.

### Transient transfections

H1299 cells were grown to approximately 70% confluency and reovirus pcDNA3 plasmids were transiently transfected with lipofectamine 2000 (ThermoFisher) at a ratio of 1:2.5 (DNA:Reagent) according to the manufacturers instructions. Forty-eight hours post-transfection cells were fixed and stained for immunofluorescence.

### DsiRNA-mediated knockdowns

For DsiRNA knock-down studies, cells were transfected with DsiRNAs (Integrated DNA Technology) directed against either an irrelevant control (G7L, vaccinia virus gene, Antisense Sequence: rUrUrUrArUrUrUrGrArUrGrArArUrCrUrArGrUrUrGrGrUrUrCrUrC Sense Sequence: rGrArArCrCrArArCrUrArGrArUrUrCrArUrCrArArArUrAAA), S2 (reovirus σ2 protein, Antisense Sequence: rCrCrUrCrUrUrArArArCrUrGrUrUrGrArGrUrCrUrGrArUrCrUrGrC Sense Sequence: rArGrArUrCrArGrArCrUrCrArArCrArGrUrUrUrArArGrAGG), or M2 (reovirus μ1 protein, Antisense Sequence: rGrArCrCrCrUrGrArGrArUrGrArArUrUrArUrUrArUrCrUTT Sense Sequence: rArArArGrArUrArArUrArArUrUrCrArUrCrUrCrArGrGrGrUrCrArG), and along with 5’ TYE™ 563 (Integrated DNA Technology) modification for immunofluorescence experiments. Cells were plated and transfected simultaneously by collecting cells via trypsinization and diluting to 2x10^5^ cells/mL and mixing with a pre-incubated cocktail of 1:2 volume Lipofectamine 2000 (11668-019, Invitrogen) to 2mM DsiRNA in Opti-MEM (31985-070, Gibco) prior to plating 12mL in 50mm^2^ dishes. Cells were allowed to adhere for at least 6 hours prior to infection. Infections and lysate collection was carried out as described above.

## ACKNOWLEDGMENTS

This publication is supported through project grants to MS from the Canadian Institutes of Health Research (CIHR), a salary award to MS from the Canada Research Chairs (CRC), infrastructure support to MS from the Canada Foundation for Innovation (CFI) and a generous donation by Linda M. Youell in memory of her husband, Gerry. J.K. received scholarships from the University of Alberta Faculty of Medicine and Dentistry 75^th^ anniversary award, a Li Ka Shing Institute of Virology Entrance Award and an Alberta Graduate Excellence Scholarship. We thank the laboratory of Dr. Gotte for the generous use of their Typhoon phosphorimager, and the University of Alberta Cell Imaging Facility for access to outstanding microscopes.

---

**Supplemental Figure S1.**
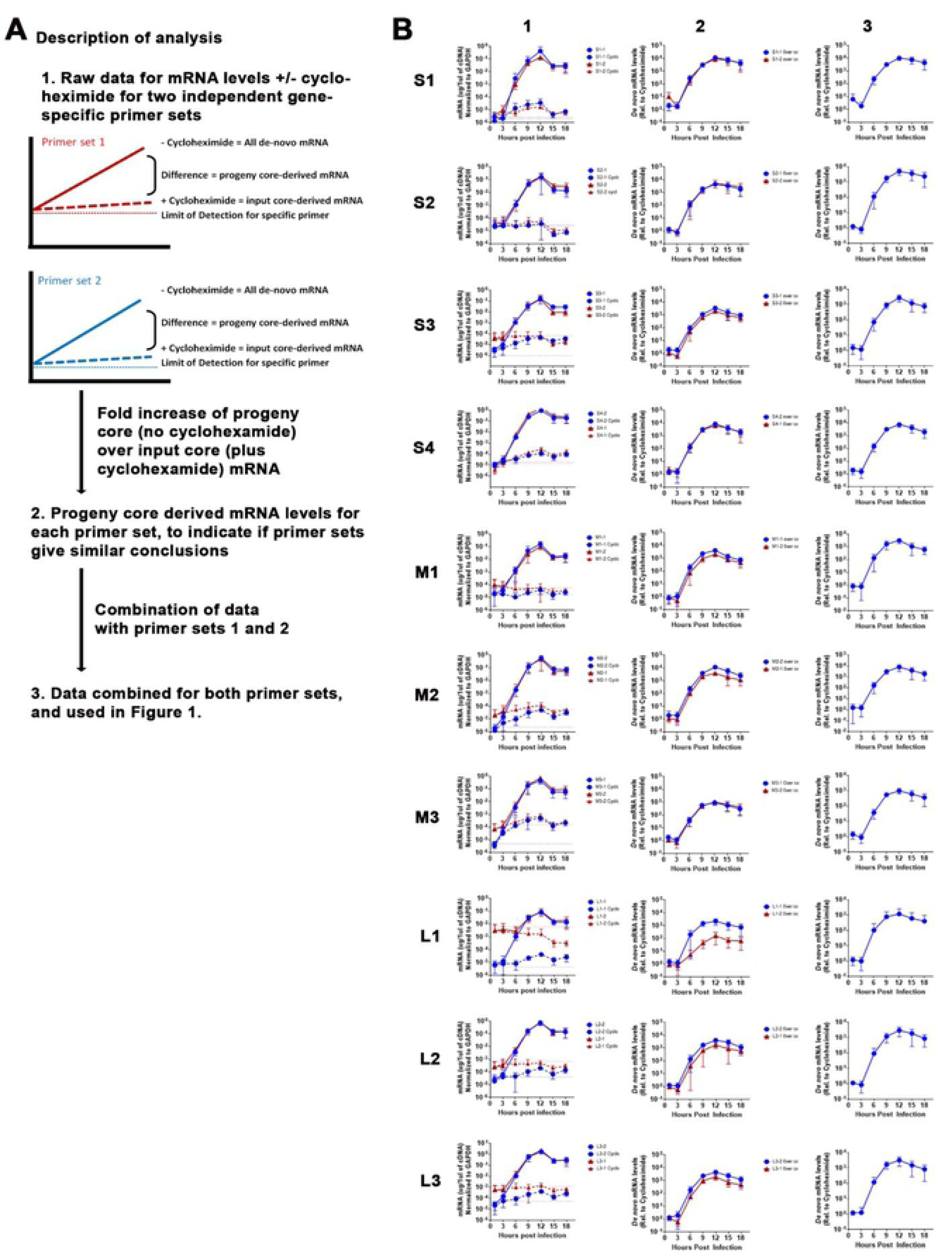
L929 cells were infected with reovirus at an MOI of 3 with or without cycloheximide (100μg/mL) and samples were collected every 3 hours to measure mRNA levels by RT-qPCR of each reovirus gene. **(A)** Description of how data was collected and analyzed. **(B-1)** Gene expression levels as measured by RT-qPCR using 2 independent primer sets, in the presence or absence of cycloheximide. **(B-2)** Difference in expression levels over cycloheximide measured by 2 independent primer sets. **(B-3)** Combined gene expression levels using two independent primer sets.

**Supplemental Figure S2.**
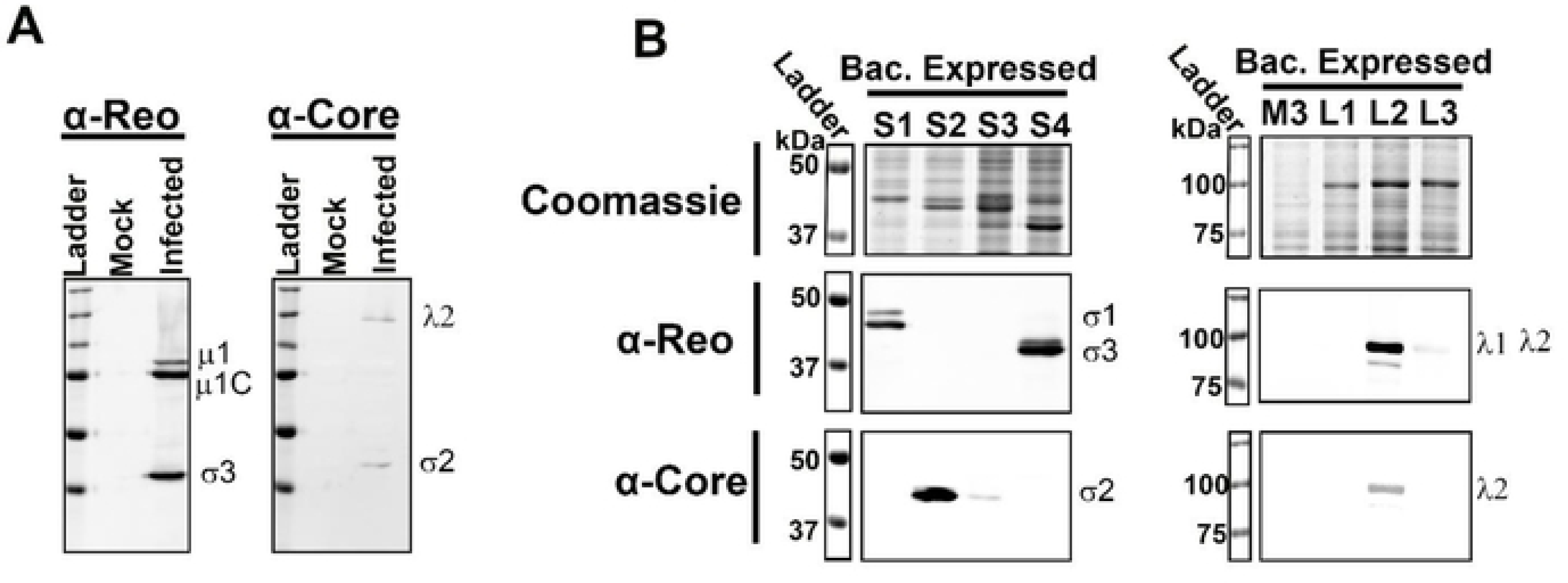
Reovirus polyclonal antibodies recognize structural proteins. **(A)** L929 cells were infected with reovirus at an MOI of 3, collected and lysed at 14 hpi. Mock-infected and infected lysates were subject to SDS-PAGE and Western blot analysis using polyclonal antibodies raised against whole virus (α-Rco, left) or (α-Core, right). **(B)** Sf9 insect cells were infected with baculoviruses expressing reovirus proteins. Cells were lysed and proteins were analyzed by SDS-PAGE and coomassie staining (top) or Western blot (middle, α-Rco and bottom, α-Core).

**Supplemental Figure S3.**
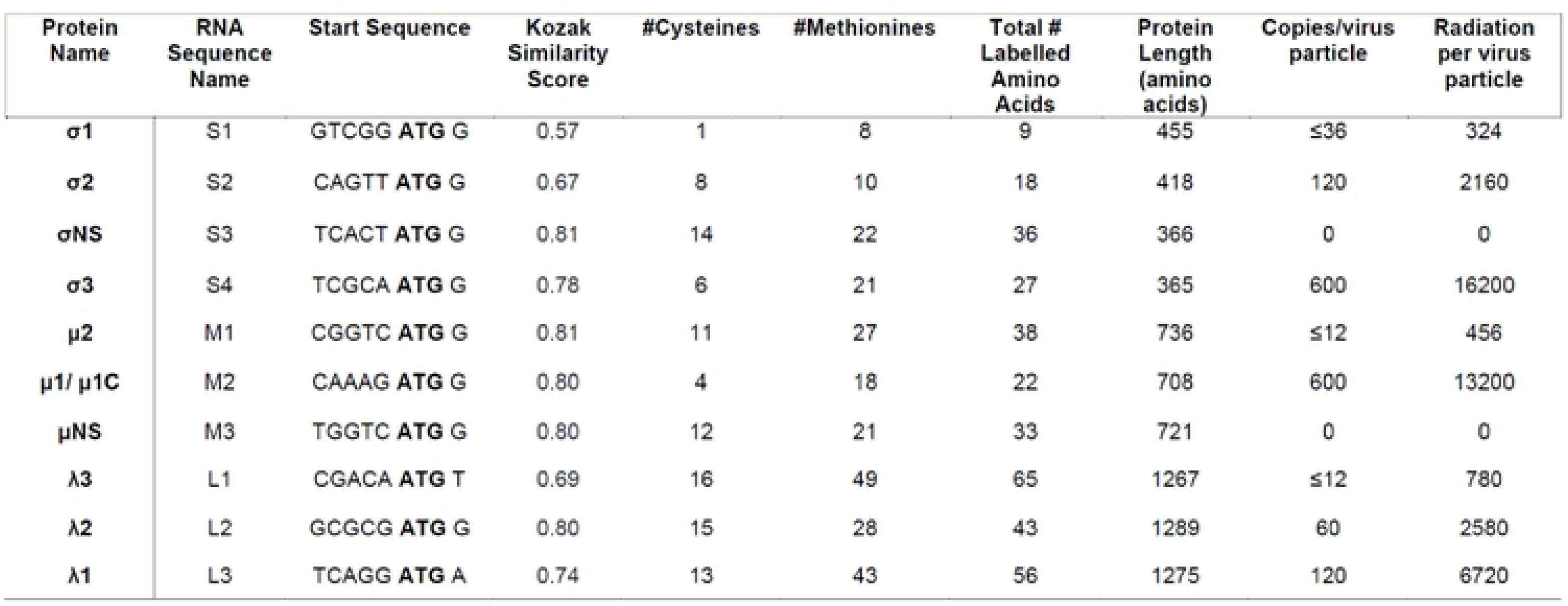
Characteristics of reovirus genome segments and corresponding proteins. RNA translation efficiency measured by Kozak similarity score and the number of labelled amino acids through incorporation of S^35^ labelled cysteine and methionine. The relative levels of radiation per protein per virus particle are also indicated.

**Supplemental Figure S4.**
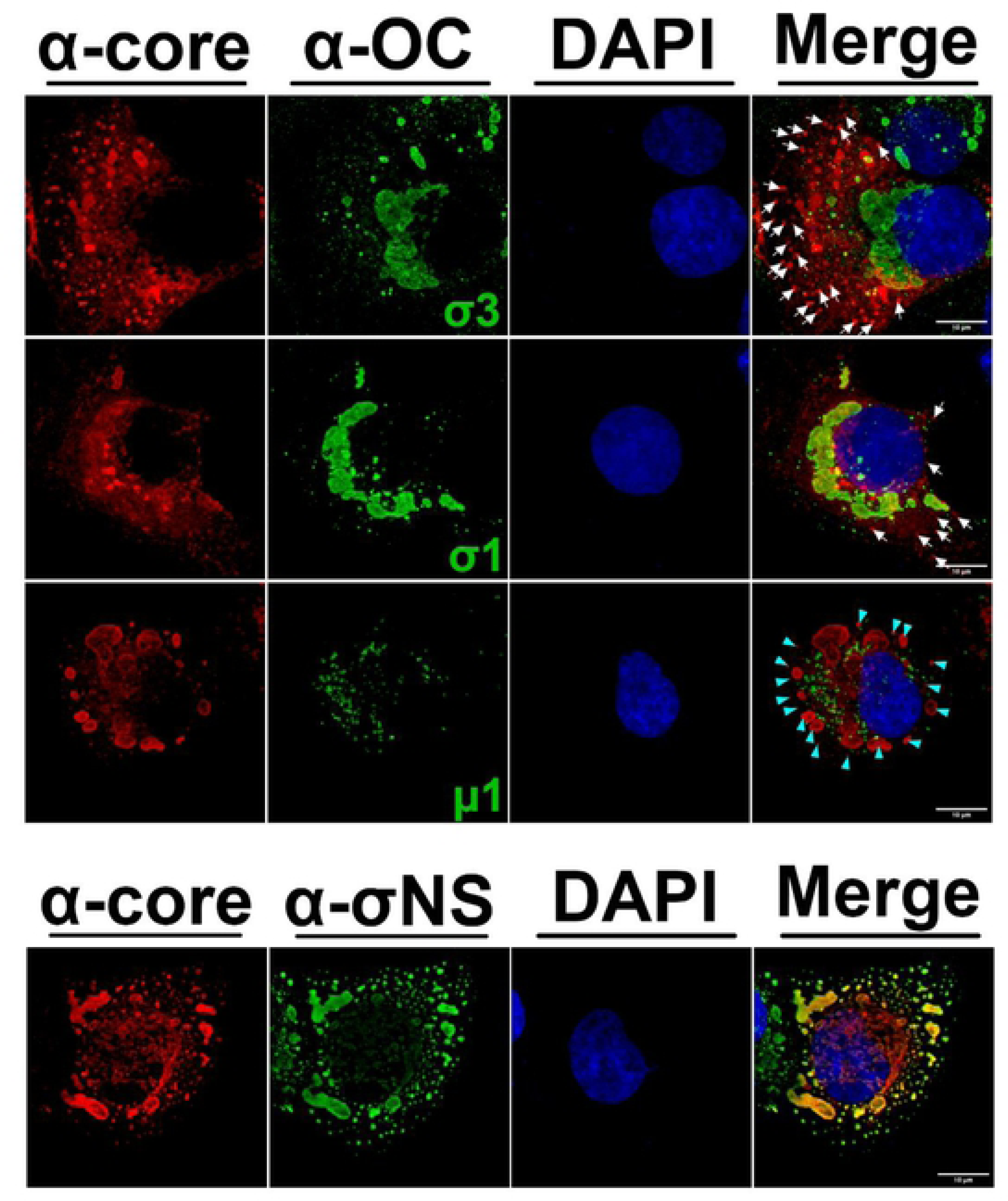
Reovirus proteins are spatially compartmentalized in T47D cells. T47D cells were infected with reovirus at an MOI of 3 before fixation at 22 hpi. Immunofluorescence staining was conducted with antibodies specific to OC proteins indicated in green (monoclonal 10G10 for σ3, monoclonal G5 for σl, or monoclonal 10F6 for μl as indicated) or σNS (monoclonal 2A9, bottom). The OC proteins were detected with secondary antibodies conjugated to Alexa 488 (pseudo colored green) or Alexa 647 (pseudo-colored red). Co-immunofluorescence in the same cells was conducted using polyclonal rabbit antibodies raised against reovirus cores (α-Core) detected with secondary antibodies conjugated to Alexa 647 (red). In the merged images, white arrows show example regions of core-only staining, while cyan arrows indicate regions of core-positive but μl-negative. Similar results were also obtained with monoclonal 10OC1 and F42 for σ3, 8H6 for μ1, and rabbit σ1-specific polyclonal serum (data not shown).

**Supplemental Figure S5.**
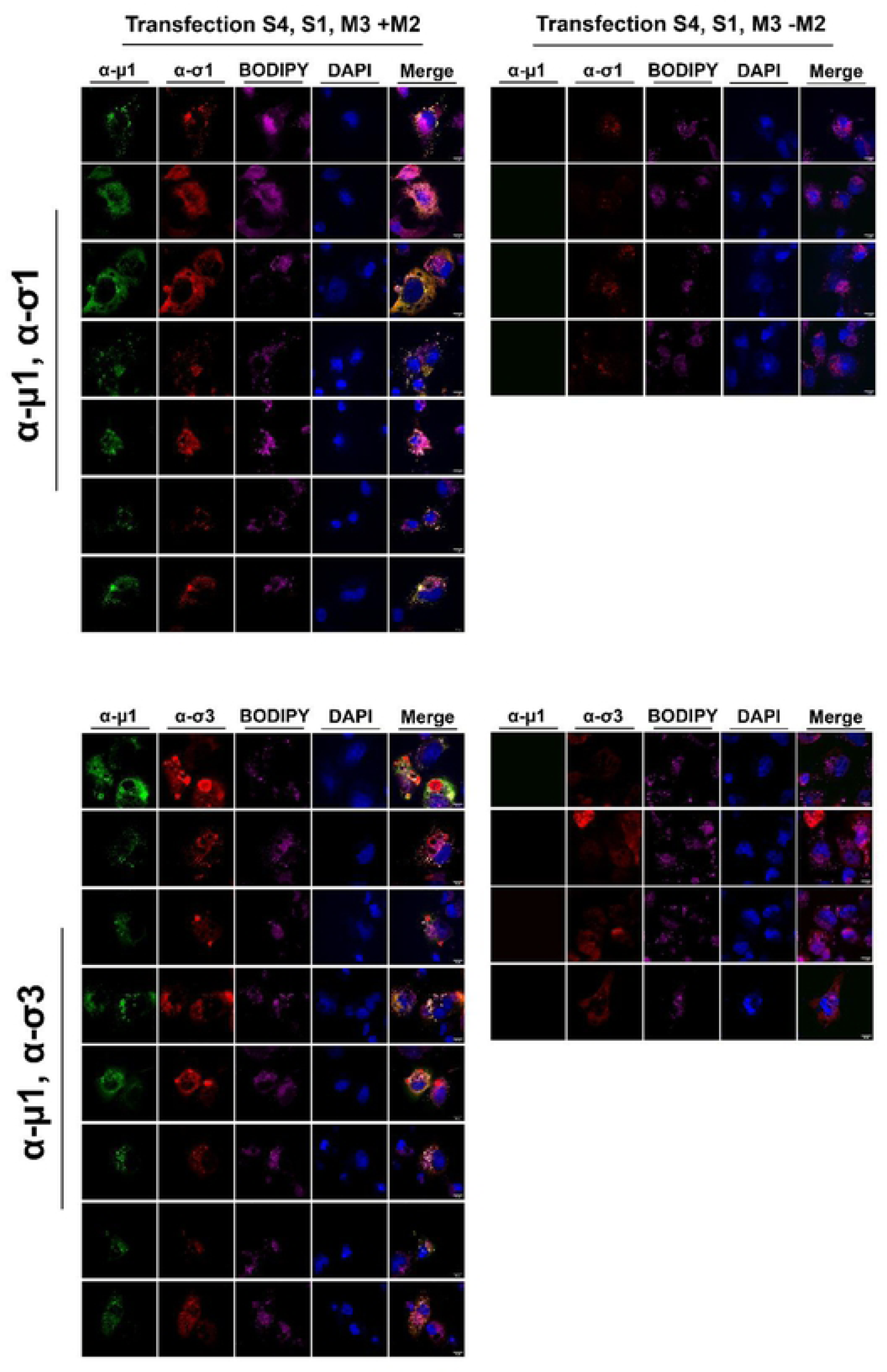
Reovirus outercapsid proteins localize to lipid droplets and perinuclear regions. H1299 cells were transfected with SlpcDNA3 (σl), S4pcDNA3 (σ3), and M3pcDNA3 (μNS) with (left) or without M2pcDNA3 (μ1) (right). Immunofluorescence staining was conducted with antibodies specific to OC proteins μl (monoclonal 10F6) and σ1 (monoclonal G5 directly labelled with AlexaFluor 647) (top) or μ1 (10F6) and σ3 (monoclonal 10C1 directly labelled with AlexaFluor 647) (bottom), BODIPY 568 for LDs, and DAPI for nuclei, μl was detected with secondary’ antibodies conjugated to AlexaFluor 488. Represented images were created from Z-stacks acquired using immunofluorescent spinning disk confocal microscopy.

**Supplemental Figure S6.**
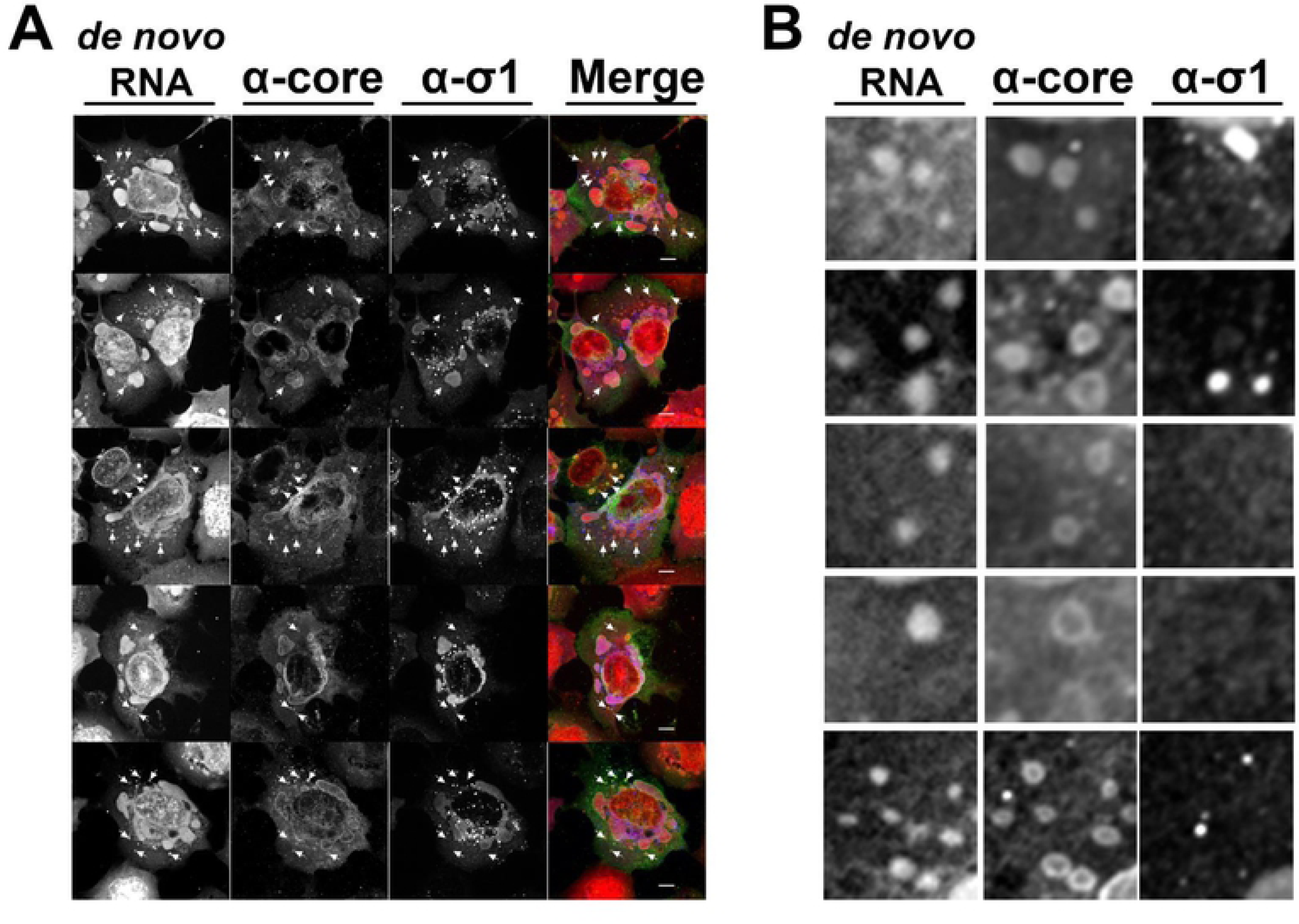
Core-containing factories are transcriptionally active. **(A)** H1299 cells were infected with reovirus at an MOI of 3 and at I4hpi, cells were treated with actinomycin D to reduce cell transcription. Between 15 hpi and 18 hpi, cells were stained for de novo transcribed RNA using an EZ-click RNA Labelling kit (RNA, red in merged image). Fixed cells were processed for immunofluorescence with rabbit polyclonal α-core antibodies (Alexa Fluor 488, green in merged image) and monoclonal mouse α-σl antibody G5 (Alexa Fluor 405, blue in merged images). Images were captured by spinning disk confocal microscopy. Scale bars represent 20μm. White arrows represent example regions positive for core and RNA staining, but negative for σl. **(B)** Close-up images show examples of OC negative, core­positive foci, which are consistently also positive for de novo RNA.

**Supplemental Figure S7.**
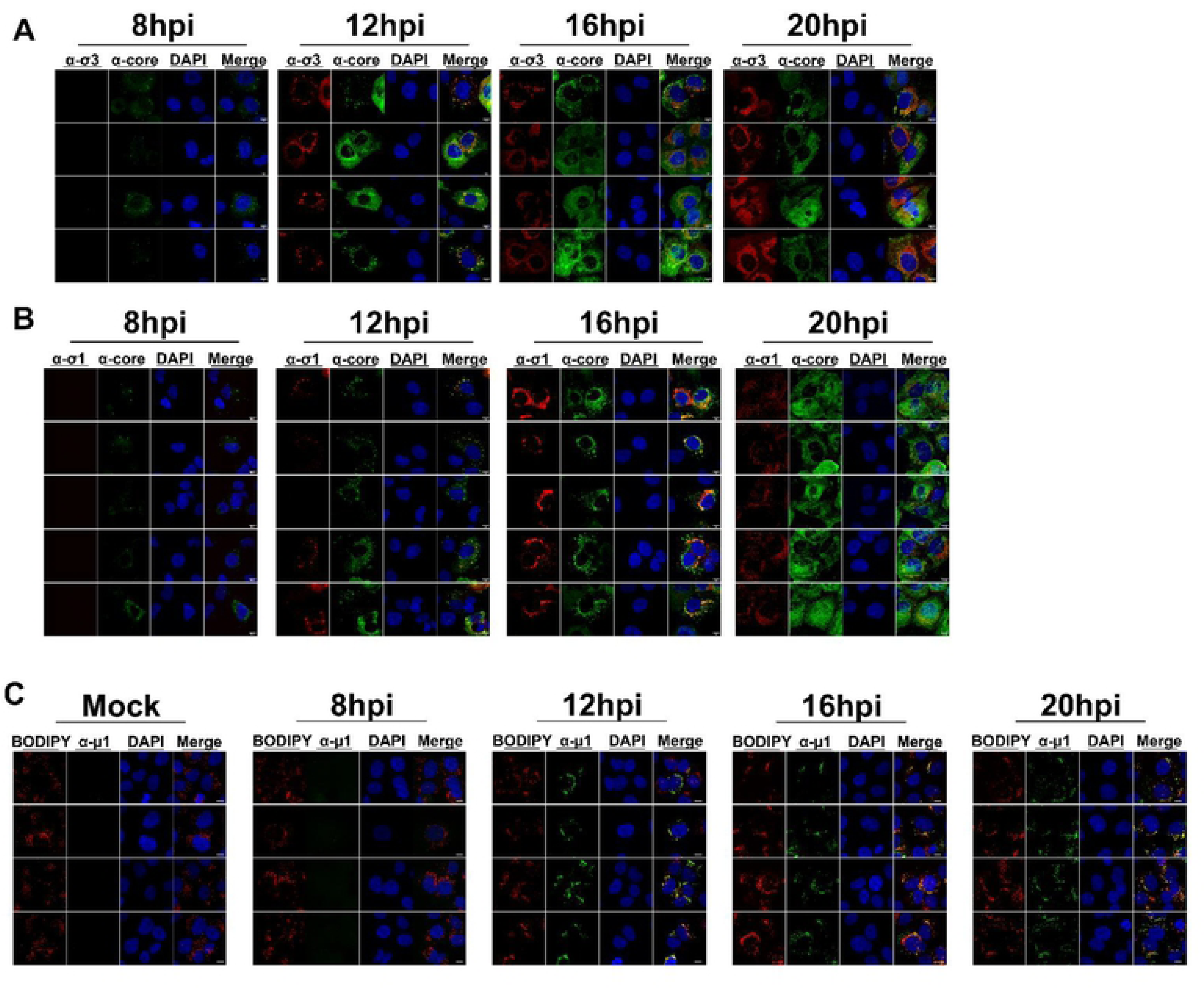
Temporal and spatial changes to reovirus compartmentalization occur over the course of infection. Representative images of H1299 cells that were infected with reovirus at an MOI of 3 and fixed at the indicated timepoints. **(A)** Immunofluorescence spinning disk confocal microscopy with monoclonal mouse α-σ3 (10G10, Alexa Fluor 647, red) and polyclonal rabbit α-core (Alexa Fluor 488, green). Cell nuclei were stained with DAPI (blue). **(B)** Immunofluorescence spinning disk confocal microscopy with monoclonal mouse α-σ1 (G5, Alcxa Fluor 647, red) and polyclonal rabbit α-core (Alexa Fluor 488, green). Cell nuclei were stained with DAPI (blue). (C) Immunofluorescence spinning disk confocal microscopy with monoclonal mouse α-μ1 (10F6, Alcxa Fluor 488, green) and BODIPY (lipid droplets, red). Cell nuclei were stained with DAPI (blue).

**Supplemental Figure S8.**
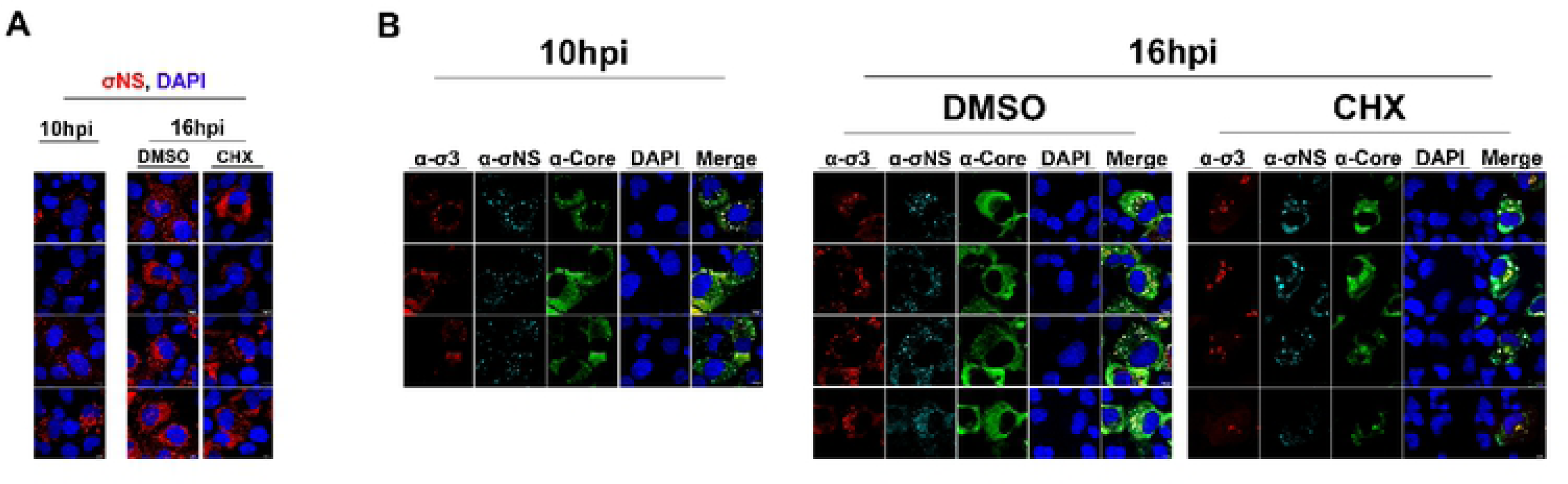
Peripheral viral factories are diminished and diffuse if translation is inhibited during early infection. **(A, B)** Representative images of Hl 299 cells were infected with reovirus at an MOI of 3, and at IO hpi cells were either fixed (A, B **left)** or treated with K)0μg/mL cyclohcximidc or DMSO (B **right).** (A) Cells were immunofluorcsccntly labelled with monoclonal mouse α-3E10 (σNS directly conjugated to AlcxaFluor 568, red) in combination with DAPI for nuclei staining (blue). (B) Cells were fixed and immunofluorescently labelled with monoclonal mouse α-σ3 (10G10, AlexaFluor 647, red), monoclonal mouse α-3E10 (σNS directly conjugated to AlexaFluor 568, cyan), polyclonal rabbit α-core (Alexa Fluor 488, green) and DAPI for nuclei staining (blue). All images were acquired via spinning disk confocal microscopy.

**Supplemental Figure S9.**
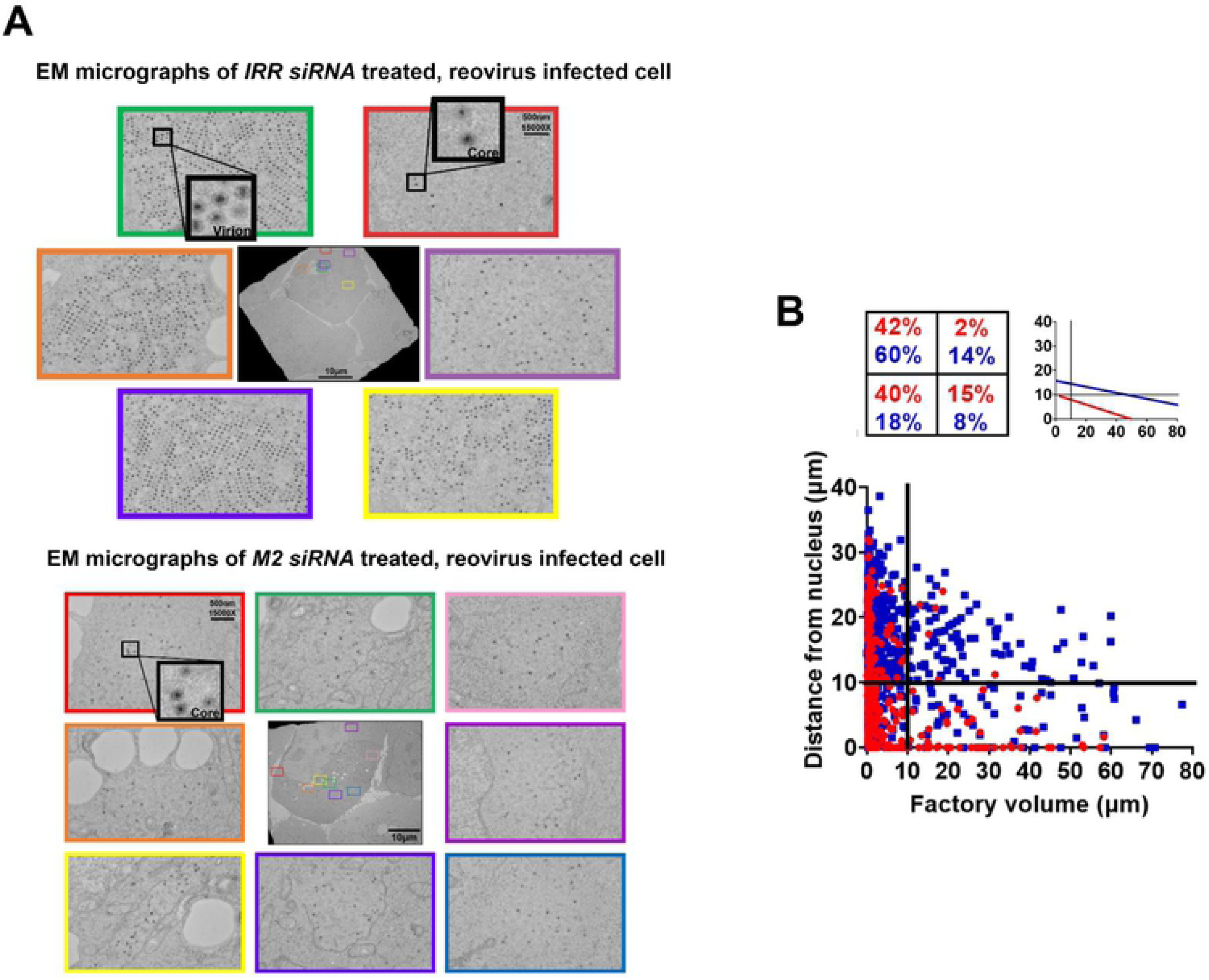
Outercapsid protein μ1 promotes convergence of peripheral core particles into perinuclear regions for whole-virion generation. Prior to infecting H1299 cells with reovirus at an MOI of 3, cells were transfected with DsiRNAs: an irrelevant control DsiRNA (IRR) or test condition (M2 gene; μ1 reovirus outercapsid protein). **(A)** Cells were fixed at 17hpi and imaged by TEM. Example images from various regions around a representative irrelevant control DsiRNA (IRR, top) or M2/μl (bottom) DsiRNA-transfected cell show the particle composition of factories found in each, with core regions only found in IRR siRNA transfected cells. Select particles are highlighted in black boxes. **(B)** The volume and distance from the nucleus were measured for each factory in ΓRR (blue squares) versus M2/μ1 (red circles). DsiRNA-trcatcd cells, using α-core channel to capture both core-only and core+OC shared factories. Based on the graph (same as found in Figure 9F), quadrants were established based arbitrarily at 10μm^3^ volume and 10μm distance from the nucleus to compare ratio of factories between IRR- and M2/μl DsiRNA-treated cells. (Top left) The percent of factories in each quadrant. (Top right) Linear regression analysis to depict trends in the distribution of factories. Data represents 8 images per condition and is representative of 2 independent experiments.

**Supplemental Figure S10.**
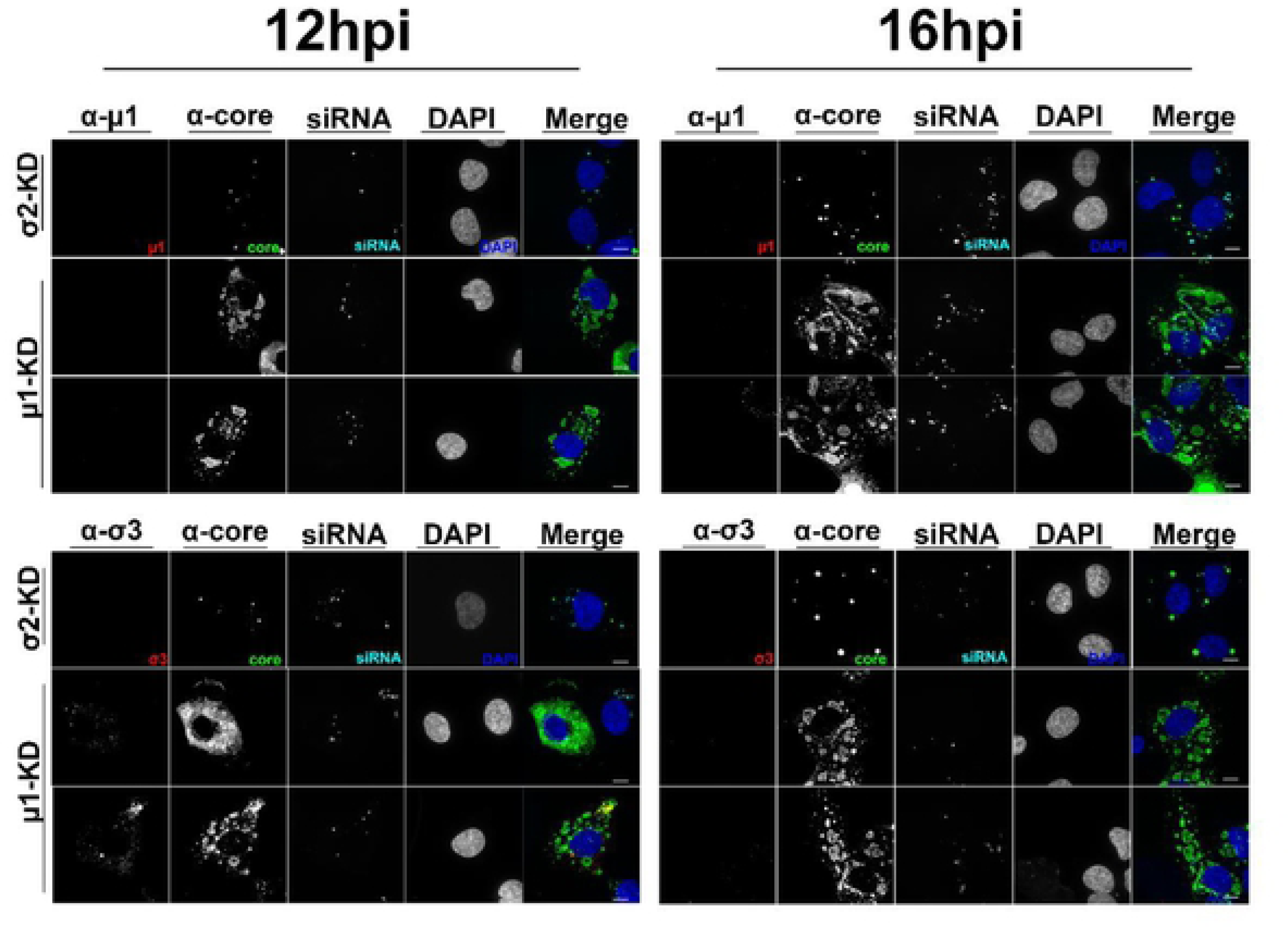
Outercapsid protein μl promotes convergence of peripheral core-only factories into perinuclear core-plus-outercapsid factories. Representative images of DsiRNA-treated infected cells at 12hpi and 16hpi. Cells were immunofluorescently labelled with monoclonal mouse α-μ1 (Top, 10F6, Alcxa Fluor 647, red in merged images) or monoclonal mouse α-σ3 (Bottom, 10G10, Alcxa Fluor 647, red in merged images) in combination with polyclonal rabbit α-core (Alexa Fluor 488, green in merged images), Tye563 for siRNA staining (cyan), and DAPI for nuclei staining (blue). All images were acquired via immunofluorescence spinning disk confocal microscopy.

**Supplemental Figure S11.**
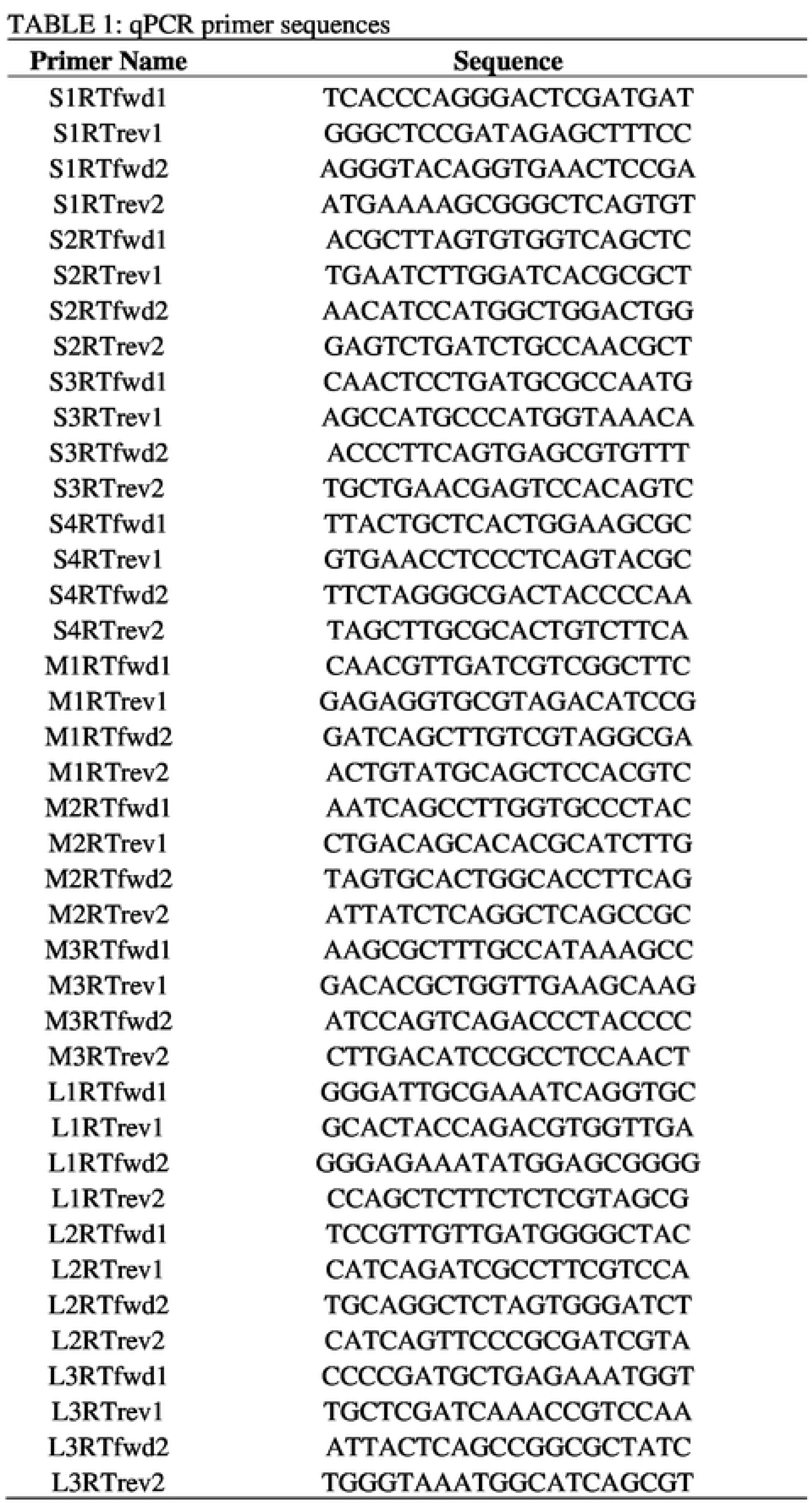
qPCR Gene Specific Primer List. RT-PCR reactions were executed following the Sybr Select (4472920, Invitrogcn) protocol using reovirus gene-specific primers listed.

